# Gene drives and population persistence vs elimination: the impact of spatial structure and inbreeding at low density

**DOI:** 10.1101/2021.11.11.468225

**Authors:** PJ Beaghton, Austin Burt

## Abstract

Synthetic gene drive constructs are being developed to control disease vectors, invasive species, and other pest species. In a well-mixed random mating population a sufficiently strong gene drive is expected to eliminate a target population, but it is not clear whether the same is true when spatial processes play a role. In species with an appropriate biology it is possible that drive-induced reductions in density might lead to increased inbreeding, reducing the efficacy of drive, eventually leading to suppression rather than elimination, regardless of how strong the drive is. To investigate this question we analyse a series of explicitly solvable stochastic models considering a range of scenarios for the relative timing of mating, reproduction, and dispersal and analyse the impact of two different types of gene drive, a Driving Y chromosome and a homing construct targeting an essential gene. We find in all cases a sufficiently strong Driving Y will go to fixation and the population will be eliminated, except in the one life history scenario (reproduction and mating in patches followed by dispersal) where low density leads to increased inbreeding, in which case the population persists indefinitely, tending to either a stable equilibrium or a limit cycle. These dynamics arise because Driving Y males have reduced mating success, particularly at low densities, due to having fewer sisters to mate with. Increased inbreeding at low densities can also prevent a homing construct from eliminating a population. For both types of drive, if there is strong inbreeding depression, then the population cannot be rescued by inbreeding and it is eliminated. These results highlight the potentially critical role that low-density-induced inbreeding and inbreeding depression (and, by extension, other sources of Allee effects) can have on the eventual impact of a gene drive on a target population.

## 1 Introduction

Gene drive is a process of biased inheritance whereby a genetic element can be transmitted from parents to offspring at a greater-than-Mendelian rate and thereby increase in frequency in a population (Burt & Crisanti, 2018). Many naturally-occurring gene drive systems have been described (Burt & Trivers, 2006; Lindholm *et al.*, 2016; Fishman & McIntosh, 2019; Burga *et al.*, 2020), and there is increasing interest in potentially using synthetic drivers to control disease vectors, harmful invasive species, and other pests (Bier, 2021; Hay *et al.*, 2021; Nolan, 2021). This interest derives in part from the fact that driving elements can spread in populations even if they cause some harm to the organisms carrying them, even disrupting reproduction to such an extent that the population could be substantially suppressed or eliminated (Burt, 2003; Godfray *et al.*, 2017). Potential strategies for population suppression include the use of gene drive constructs to produce a male-biased sex ratio, or to knock-out genes needed for survival or reproduction, or both (Galizi *et al.*, 2014; Kyrou *et al.*, 2018; Simoni *et al.*, 2020).

Because drive depends on a deviation from Mendelian transmission it cannot operate in wholly asexual populations, and, moreover, it will tend to be less effective in inbred populations (i.e., with mating of close relatives), where the frequency of heterozygotes is reduced relative to outcrossed populations (Burt & Trivers, 2006; Agren & Clark, 2018). The extent of inbreeding in a population can be affected by many factors, including, potentially, population density. In particular, in some species, when densities are low, the only mates available may be relatives, and the frequency of inbreeding correspondingly high. In such a species, release of a gene drive could lead to a reduction in population density, which in turn leads to increased inbreeding, reducing the effectiveness of the drive and the ultimate impact on population density (relative to what would have occurred had there been no change in inbreeding), potentially even making the difference between the target population persisting or being eliminated (Bull *et al.*, 2019).

Previous modelling has investigated some aspects of this problem. The reduced efficacy of drive in the face of inbreeding has been analysed in numerous contexts, including the autosomal killers (Petras, 1967), B-chromosomes (Burt & Trivers, 1998), transposable elements (Wright & Schoen, 1999), and engineered gene drive constructs for population suppression (Drury *et al.*, 2017). In each case the breeding system was treated as an exogenously determined variable. Hamilton (1967) demonstrated that, in species whose biology is such that low density leads to increased inbreeding, low density could be a barrier to the spread of a Driving Y chromosome. Again, population density in his model was an exogenous variable, rather than an endogenous one responding to the presence of the Driving Y. In the closest precedent for the modelling presented here, Bull *et al.* (2019) analysed the impact of two different types of gene drives on a population when the frequency of sib mating is assumed to increase as population mean fitness declines, and found that population elimination could be prevented, even with perfect drives. However, they did not model population density explicitly. Finally, while deterministic spatial models using partial differential equations can show population elimination by sufficiently strong drives (Beaghton *et al.*, 2016), stochastic individual-based models often lead to suppression but not elimination (North *et al.*, 2013, 2019, 2020; Eckhoff *et al.*, 2017), potentially consistent with a role for low density inbreeding, though inbreeding was not monitored or manipulated in these models. More recently, Champer *et al.* (2021) analysed an individual-based model of gene drives in continuous space, and observed that preventing inbreeding promoted elimination, consistent with expectations, but did not study this result in detail.

To more fully investigate the potential role of low-density-induced inbreeding in preventing population elimination, we have analysed a series of explicitly solvable stochastic models that include spatial structure, gene drive, and alternative life history scenarios of mating, dispersal, and reproduction. We first focus on Driving Y chromosomes, and consider seven life history scenarios. In all of them a sufficiently strong Driving Y will eliminate a population, except the one scenario in which low population density leads to increased inbreeding, in which case there is suppression but not elimination, no matter how strong the drive. We then show that the same life history also prevents population elimination by a gene drive that uses the homing reaction. In both cases populations persist because inbreeding gives a fitness advantage to the wildtype chromosome over the driver; incorporating strong inbreeding depression into the models removes this fitness advantage, and the population is then eliminated. These results highlight the key role that low-density-induced inbreeding can have on the fate of a population faced with a gene drive, and emphasize the importance of incorporating inbreeding depression (and, by extension, other negative effects of low density on population growth rates) in models of suppressive gene drives.

## 2 Driving Y

We model an infinite sized population with discrete generations. Two key events in a species’ life history are mating of males and females, and offspring production by mated females to make the (unmated) males and females of the next generation. Each of these activities can occur either in an infinite well-mixed population (“in the cloud”), or after individuals have settled randomly into an infinite array of “patches”, so in addition to mating and reproduction there is also movement. Mating is random, so that if mating occurs in the cloud then it is according to the proportion of the different types in the cloud, whereas if it occurs in patches, then it is according to the different types in the particular patch. Females mate only once in their life, and store the sperm for subsequent reproduction, whereas a male may mate multiple times, and all females get mated as long as there is at least one male in the cloud or patch. Offspring production is density-dependent, according to the Beverton & Holt (1957) model; if reproduction is occurring in the cloud then the average number of offspring produced per female depends on the density (of mated females) in the cloud, whereas if reproduction is occurring in patches then it is the local density that counts.

We first consider the release of males carrying a Driving Y chromosome engineered to be transmitted to more than 50% of the offspring (e.g., by disrupting transmission of the X chromosome (Galizi *et al.*, 2014; Fasulo *et al.*, 2020)). There are thus two types of males, those with Wildtype (W) and Driving (D) Y chromosomes and two types of mated females, those mated to a W-male and those mated to a D-male. W-mated females produce on average equal numbers of female and W-male offspring, whereas D-mated females produce on average female and D-male offspring at a ratio (1 *− m*): *m*.

We now consider a range of scenarios for the location of mating and reproduction (cloud or patches) and the timing of movement between them. Results are summarised in Figure 1.

**Figure 1:**
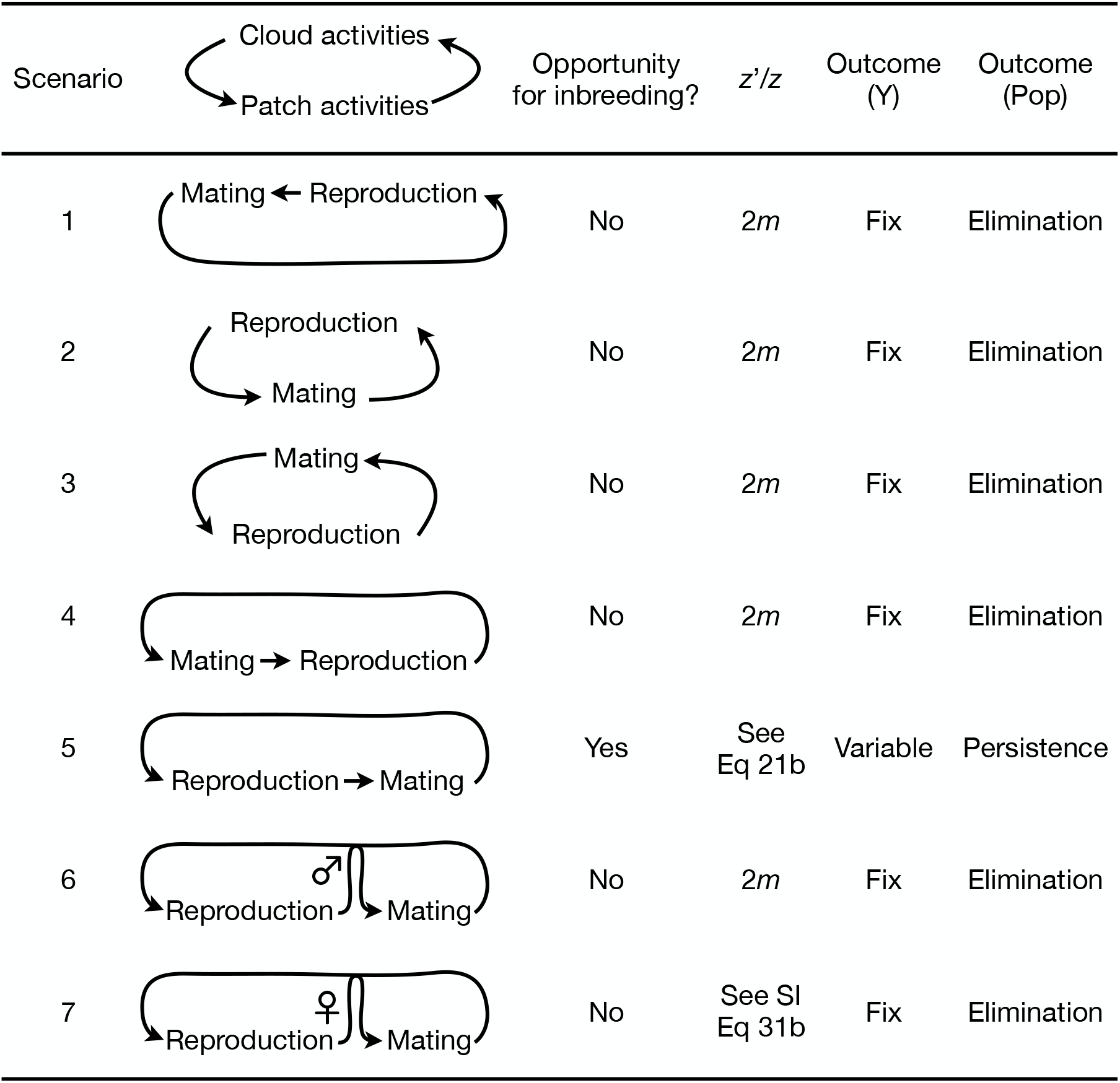
Consequences of alternative life histories for the fate of a Driving Y and the population. *z*′*/z*: Odds ratio for a male carrying a Driving Y in one generation to that in the previous generation. Outcome (Y): Outcome for the proportion of males carrying the Driving Y. Fix: Driving Y goes to fixation; Variable: Driving Y may go to fixation, remain polymorphic, or be lost. Outcome (Pop): Outcome for the population assuming a sufficiently strong drive (e.g., *m* = 1).

### 2.1 Scenario 1: A well-mixed population

Our starting point is a non-spatial model in which both mating and offspring production occur in the cloud, which is of infinite size and contains individuals at a finite density. The Driving Y is introduced at a given density at *t* = 0 into a wildtype (W) population at equilibrium. The (finite) population density then evolves from generation to generation as

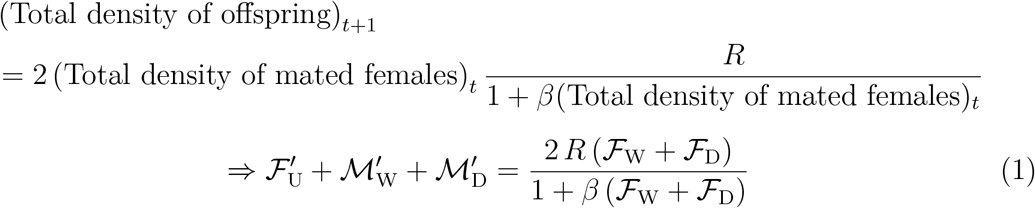

where 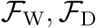 are the densities of W- and D-mated females in generation *t* and 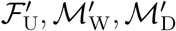 are the densities of female, W-male and D-male offspring in generation *t* + 1, *R* is the intrinsic (or low density) rate of increase of the population, and *β* is a parameter describing the strength of density dependence, where 1/*β* is the density at which the population growth rate is half its maximum value.

Before the release of the Driving Y and at equilibrium, we have 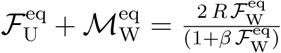. Since W-mated females produce on average equal number of female and W-male offspring and since all the female offspring turn into W-mated females, we have 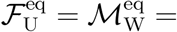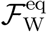. The prerelease equilibrium equation becomes

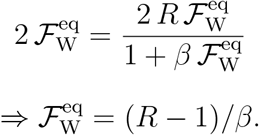

W-mated females produce female offspring and W-male offspring in equal numbers (averaged over the entire cloud) whereas D-mated females produce a skewed ratio of (1 – *m*): *m* female offspring vs D-male offspring, so

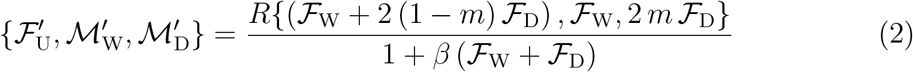

As mating is random, the probability that a female offspring becomes a W-mated female is 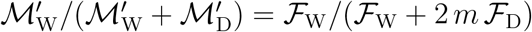 whereas the probability that she becomes a D-mated female is 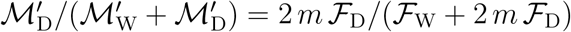. So, the densities of W- and D-mated females in generation *t* + 1 are

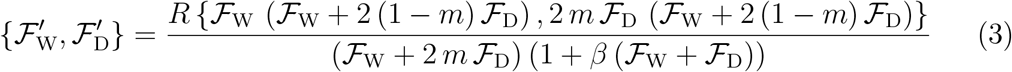

We now introduce a change of variables 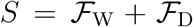 and 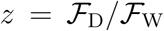 and the equations above become

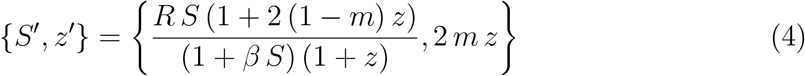

The recurrence equations (4) for {*S, z*} are sufficient to update the system from generation *t* to generation *t* + 1. Note that as long as *m* > 0.5, *z* will increase without bound, implying that the Driving Y tends to fixation and, if *m* is sufficiently large, then the population will tend to elimination – most obviously if *m* = 1, then the population will tend to be all male.

### 2.2 Scenario 2: Local mating

Now suppose male and female offspring settle randomly into patches, there is local competition within each patch among males to mate with the females, and then the mated females return to the cloud and reproduce in a density-dependent manner as described in Scenario 1. In Scenario 1, the pre-release equilibrium density of W-mated females was shown to be 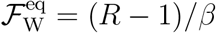.

In this model, the total density of offspring is again given by (1). We allocate the well-mixed offspring population in the cloud into (an infinite number of) patches of volume *V*. The actual (integer) number of offspring in each patch is Poisson distributed with a mean

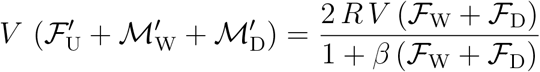

and ranges from zero to infinity. We restrict mating only among members of the same patch and all females mate if there is at least one male in the patch. Due to stochasticity, a fraction of patches will contain zero males and as a result a fraction of the females will not mate (the lower the population density in the cloud, the larger the fraction of females that will be unsuccessful in mating due to a lack of males in their patch).

Equation (2) that gives the number of offspring in the next generation holds here too, so a patch of males and females of volume *V* consists (on average) of 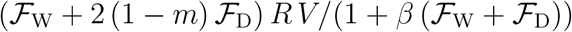 females, 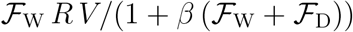 W-males and 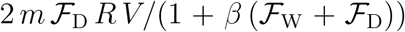 D-males. The probabilities of having a Poisson-distributed set of {F_U_, M_W_, M_D_} offspring in a patch are thus

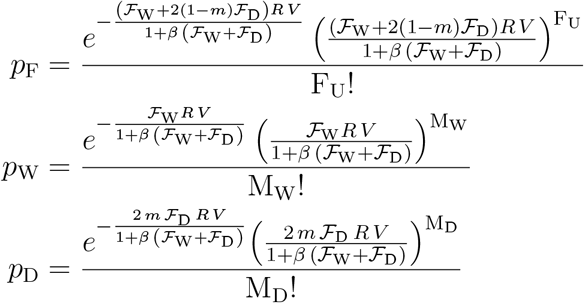

The probability that a patch contains M_W_ = M_D_ = 0 males is *p*_W_ |M_W_=M_D_=0 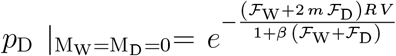, which goes to 0 when *V* → ∞ and goes to 1 when *V* → 0 (i.e. when the volume *V* is so infinitesimally small that it is certain that any patch with a female will contain no males).

The probability of *k* females, out of the F_U_ females in a patch, becoming W-mated females is 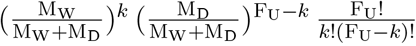 since every female undergoes a Bernoulli trial in picking a male out of the M_W_ W-males and M_D_ D-males in her patch. The expected number of W-mated females in a patch, conditional on {F_U_, M_W_, M_D_}, is thus

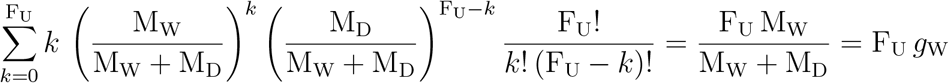

where 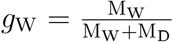 is the fraction of W-males in the patch. Similarly, the expected number of D-mated females in a patch, conditional on {F_U_, M_W_, M_D_}, is 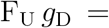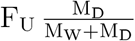.

We introduce M = M_W_ + M_D_ and use Wolfram Mathematica to evaluate the densities of W-mated and D-mated females arising from the two types of pairings, averaged over all the mating cohorts (the division by *V* converts the expected number of mated females in the mating cohorts to a density):

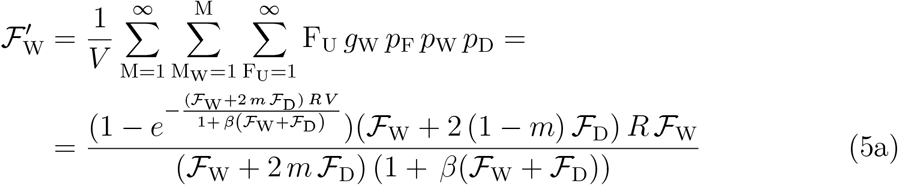

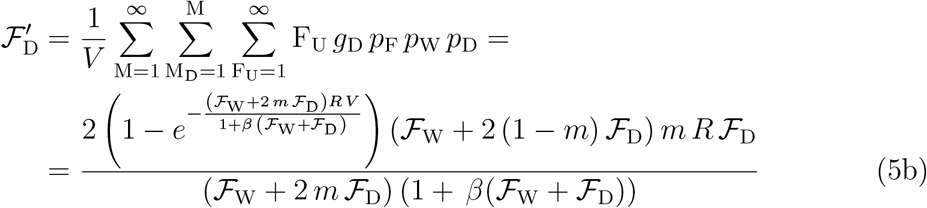

We introduce the change of variables 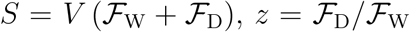 and *α* = *β/V*; equations (5a)-(5b) now become:

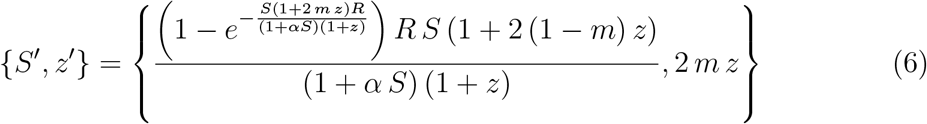

The recurrence equations (6) are again sufficient to update this system from generation *t* to generation *t* + 1. The transition equation for *z* is the same as in Scenario 1, indicating, again, that the Driving Y will go to fixation, and, if *m* is sufficiently large, the population will be eliminated.

### 2.3 Scenario 3: Local density-dependent reproduction

We now reverse the location of events, so mating occurs in the cloud and reproduction occurs in patches (subject to local density dependence). The cloud densities of W- and D-mated females in generation *t* are 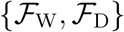, and a Poisson-distributed random sample of {F_W_, F_D_} W- and D-mated females with means 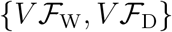, is drawn from the cloud and settles in each patch. The probabilities of having F_W_ and F_D_ mated females in a patch are

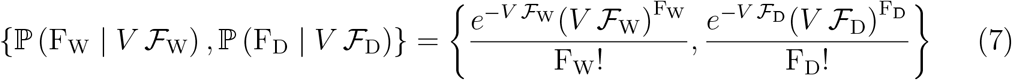

In this model the reproduction rate of mated females depends on the number of the mated females in the local patch. We assume the number of surviving offspring that each mated female produces is Poisson-distributed with a mean λ = 2 *R*/ (1 + *α* (F_W_ + F_D_)), where *α* > 0 is a density dependence parameter appropriate for patches instead of the cloud. Note that the maximum low-density rate of increase is now *R/* (1 + *α*). The probability that the F_W_ + F_D_ mated females generate *j* offspring in total in a given patch is then

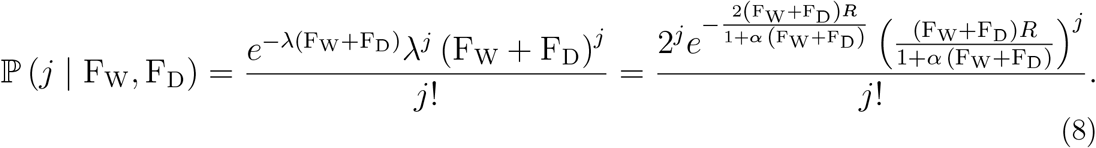

These *j* offspring are made up of *i*_F_ females, *i*_W_ W-males and *j* – *i*_F_ – *i*_W_ D-males. The probability of a {*i*_F_, *i*_W_, *j* – *i*_F_ – *i*_W_} triplet is derived from a multinomial distribution with *j* trials and normalised weights 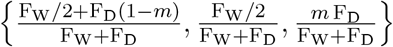, since, on average, fractions of 1/2 and 1 – *m* of W-mated and D-mated females’ offspring are female, with the rest of the offspring being W- and D-males, respectively.

Hence, the probability ℙ(*i*_F_, *i*_W_, *j* – *i*_F_ – *i*_W_ | *j*, F_W_, F_D_) of having {*i*_F_, *i*_W_, *j* – *i*_F_ – *i*_W_} female, W-male and D-male offspring in the patch (conditional on *j* total offspring from {F_W_, F_D_} mated females) using the weights above is

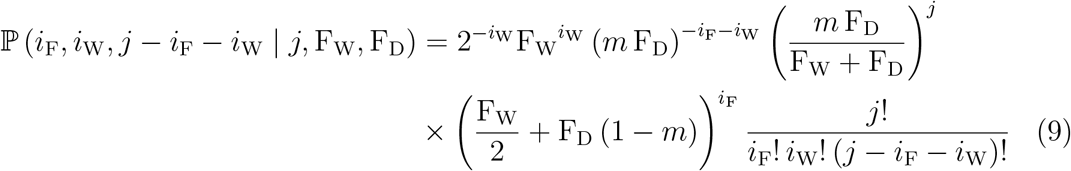

and the expected numbers of female, W-male and D-male offspring in the patch (conditional on *j* total offspring from {F_W_, F_D_} mated females) is obtained by summing over all possible values of *i*_F_ and *i*_W_:

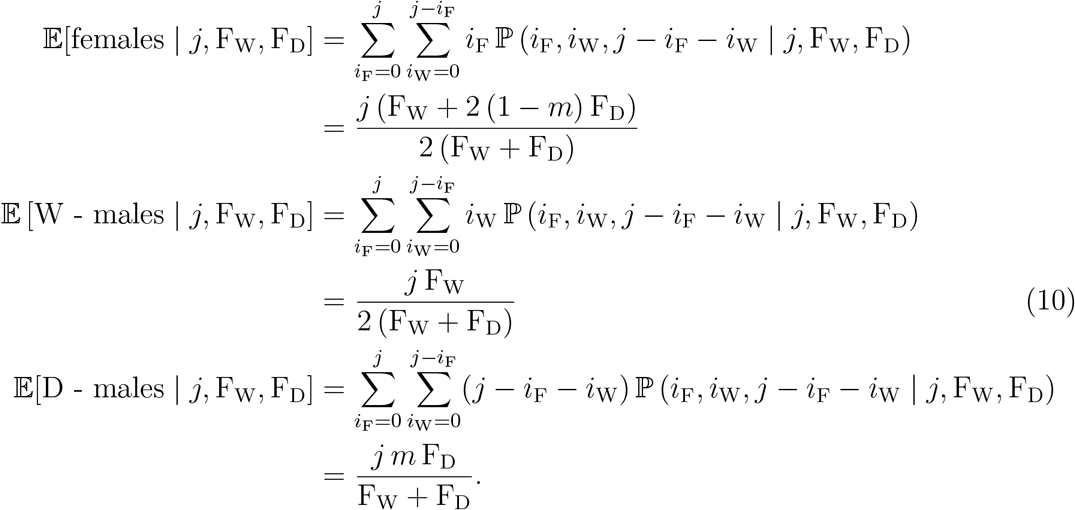

We use (7), (8) and (10) to evaluate the densities of the offspring that will aggregate back in the cloud, 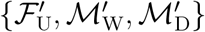, by summing the respective products over all possible values of F_W_, F_D_ and *j* from 0 to infinity and dividing by *V*:

The cloud density 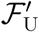 of female offspring before mating is:

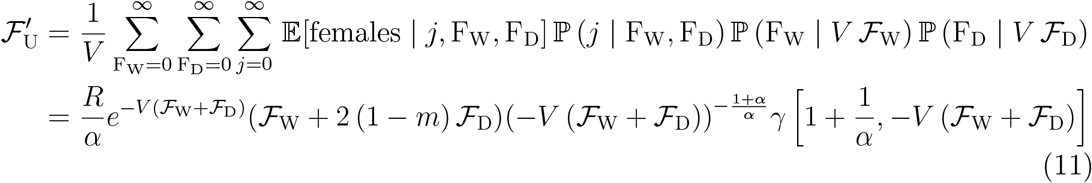

where *γ* is the lower incomplete gamma function.

The cloud density 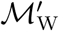 of W-male offspring is:

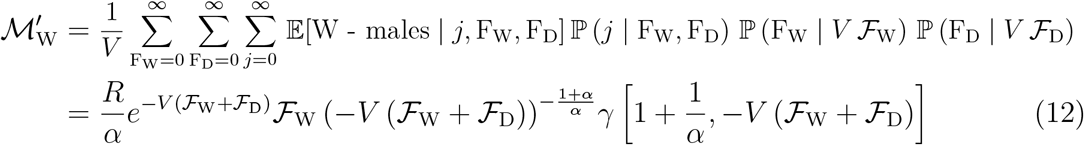

The cloud density 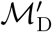 of D-male offspring is:

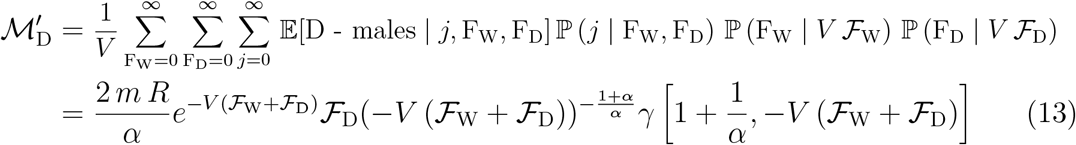

All the aggregated offspring in the cloud form a single mating pool, with each female choosing a random mate. Given that there will always be at least one male in the (infinite) mating pool, all unmated females become mated females (i.e. 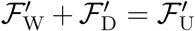) and the fractions of the resulting W- and D-mated females are simply equal to the fractions of W- and D-males in the cloud, i.e. 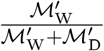 and 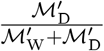. From (11),(12) and (13) it follows that the fractions of W- and D-males in the cloud mating pool reduce to 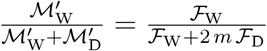 and 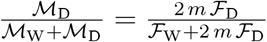, respectively. Thus, the densities of mated females in the cloud in generation *t* + 1 are

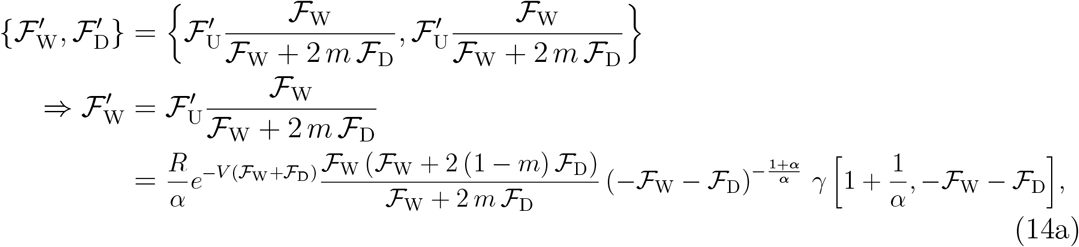

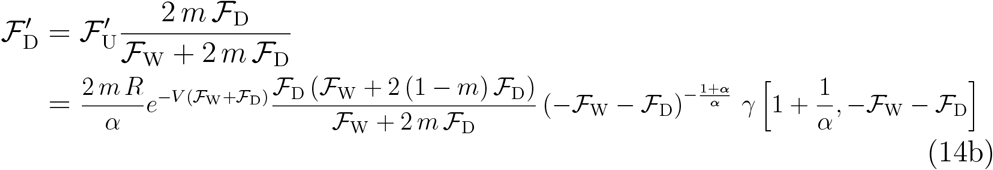

The change of variables used in previous models, 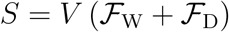 and 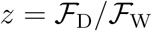, gives the recurrence equations that update the state variables from generation *t* to generation *t* + 1:

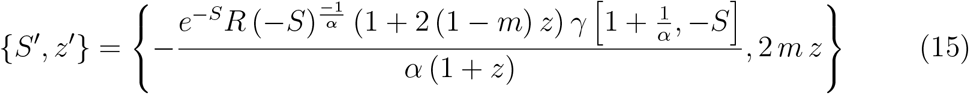

Again, the Driving Y tends to fixation, and, for sufficiently large *m*, the population will tend to elimination.

### 2.4 Scenario 4: Local mating followed by local reproduction

In this scenario males and unmated females settle randomly into patches, mate locally, reproduce in a (locally) density-dependent manner, then males and unmated females rise back again to the cloud to be re-assorted back to patches. It is convenient to derive the recurrence equations for this model by starting with the stage in generation *t* at which all the female and male offspring find themselves well-mixed in a cloud, which contains an (infinite) number of (unmated) females, W- and D-males with (finite) densities 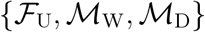. All the cloud inhabitants then settle into patches, with each patch containing *on average* 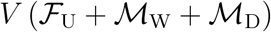 individuals. As in Scenario 2, the actual number of individuals in a patch is Poisson-distributed with means 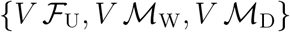, for the females, W-males and D-males, respectively. The probabilities {*p*_F_, *p*_W_, *p*_D_} of having a set of {F_U_, M_W_, M_D_} individuals in a patch are

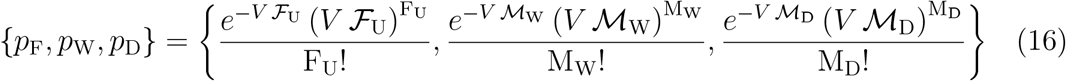

Each of the F_U_ unmated females settling in a patch will randomly chose a single W-male or D-male partner from within her patch. The probability of the F_U_ unmated females in a patch becoming {F_W_, F_U_ – F_W_} W- and D-mated females, respectively, is thus

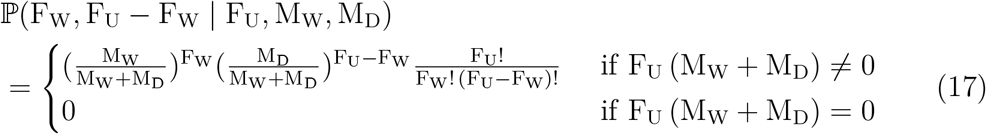

We set F_D_ = F_U_ – F_W_ and then use (8) for the probability ℙ(*j* | F_W_, F_U_ – F_W_) that the F_U_ mated females in a given patch generate *j* offspring in total and (9) for the probability ℙ (*j*– *i*_M_, *i*_W_, *i*_M_ – *i*_W_ | *j*, F_W_, F_U_ – F_W_) of having {*i*_F_ = *j* – *i*_M_, *i*_W_, *i*_D_ = *i*_M_ – *i*_W_} female, W-male and D-male offspring in the patch (conditional on *j* total offspring from {F_W_, F_U_ – F_W_} mated females).

We can now derive the new cloud densities 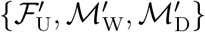, having started from 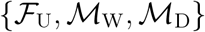 in the previous generation. We start with {*i*_F_ = *j* – *i*_M_, *i*_W_, *i*_D_ = *i*_M_ – *i*_W_} new female, W-male and D-male offspring in a patch, conditional on {F_U_, M_W_, M_D_ = M – M_W_} offspring from the previous generation having settled in the patch and having produced {F_W_, F_U_ – F_W_} mated females who in turn have produced *j* = *i*_F_ + *i*_W_ + *i*_D_ offspring in total. We then combine the various probabilities in (8), (9), (16) and (17), introduce *i*_F_ = *j* – *i*_M_ and M_D_ + M_W_ = M, and sum over all possible values of {*i*_W_, *i*_M_, *j*, F_W_, M_W_, M, F_U_} to evaluate the average numbers of female, W-male and D-male offspring (across all patches) and then divide them by *V* to convert them into the densities 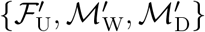 in the next generation (we use Wolfram Mathematica to evaluate each of the 7-deep nested sums):

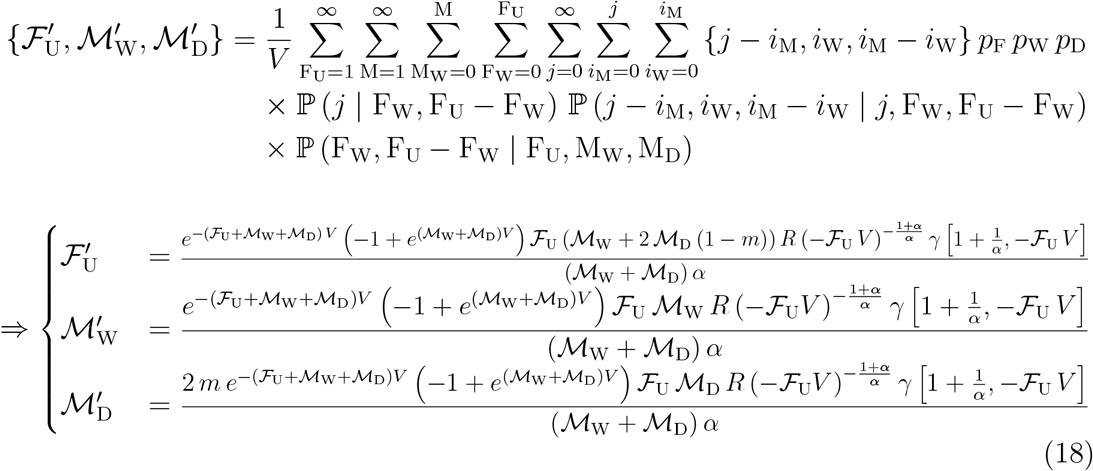

The ratio 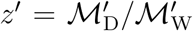 of D-male density to W-male density in the cloud can be obtained by dividing the two male densities in (18):

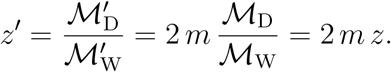

As in earlier scenaria, given that 2 *m* > 1, *z* → ∞ as *t* → ∞ and the Driving Y asymptotically fixes in the population with 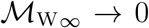 and 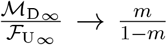, and, for sufficiently large *m*, the population will be eliminated.

### 2.5 Scenario 5: Local reproduction followed by local mating

Now suppose both mating and reproduction are again local, but the order of events is changed, such that it is mated females that disperse, rather than unmated males and females. Note that in this scenario where reproduction is local and there is no pre-mating dispersal there is the opportunity for inbreeding to occur (i.e., a female to mate with her brother), and the probability of this occurring will tend to increase as the population density decreases.

Similarly to previous scenaria, the cloud densities of W- and D-mated females in generation *t* are 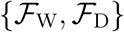, and the numbers {F_W_, F_D_} of W- and D-mated females settling in a patch is Poisson-distributed with means 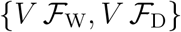.

The *j* offspring of the {F_W_, F_D_} mated females in each patch mate locally, and to obtain the expected number of new W-mated females in the patch, conditional on {*j*, F_W_, F_D_}, we use (9) from Scenario 3 for the probability ℙ (*i*_F_, *i*_W_, *j* – *i*_F_ – *i*_W_ | *j*, F_W_, F_D_) of having {*i*_F_, *i*_W_, *j* – *i*_F_ – *i*_W_} female, W-male and D-male offspring in the patch and sum over all possible *i*_F_ and *i*_W_, noting that *i*_F_ = 0 and *i*_F_ = *j* are excluded from the *i*_F_-summation as they both result in no mated females (because of either no female offspring, i.e. *i*_F_ = 0, or all female offspring, *i*_F_ = *j*, and thus no males to mate with):

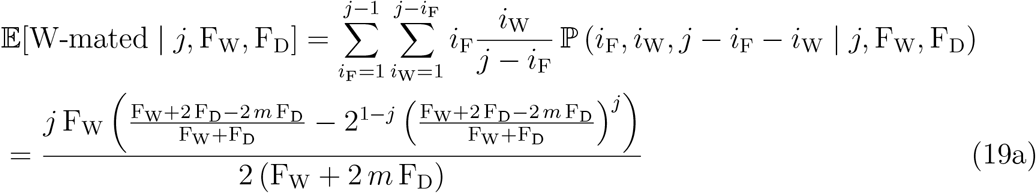

Similar analysis gives the expected number of D-mated females in each patch, conditional on *j* total offspring from {F_W_, F_D_} mated females:

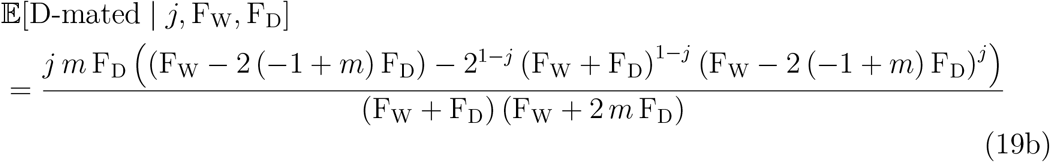

At this stage, the mated females in each patch migrate to the cloud. In order to calculate the density of W-mated females in the cloud, we combine (7) for the probabilities 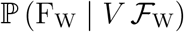 and 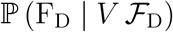 of having F_W_ and F_D_ mated females in a patch, (8) for the probability ℙ (*j* | F_W_, F_D_) that the F_W_ + F_D_ mated females generate *j* offspring in total in the patch (since scenario 3 and 5 share the same (local) density dependent reproduction mechanism) and (19a) for the expected number of W-mated females, conditional on *j* offspring from F_W_ and F_D_ mated females in the patch. Their product is then summed over all possible values of F_W_, F_D_ and *j* to give the average number of W-mated females across all patches, and is then divided by *V*, to give the expression for 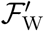, the density of W-mated females in the cloud in generation *t* + 1:

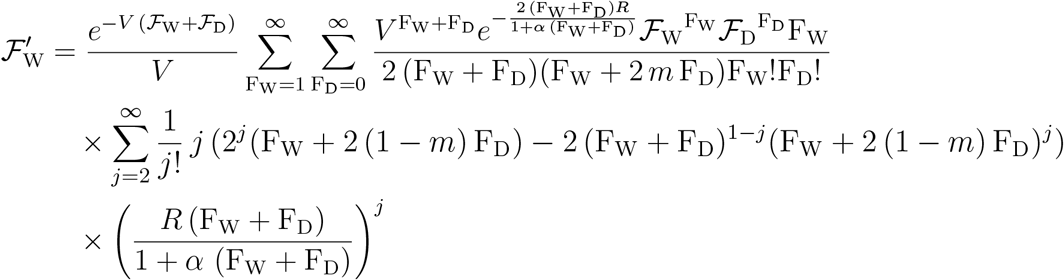

The innermost summation over *j* can be calculated analytically so the expression above reduces to a double infinite sum over F_W_ and F_D_:

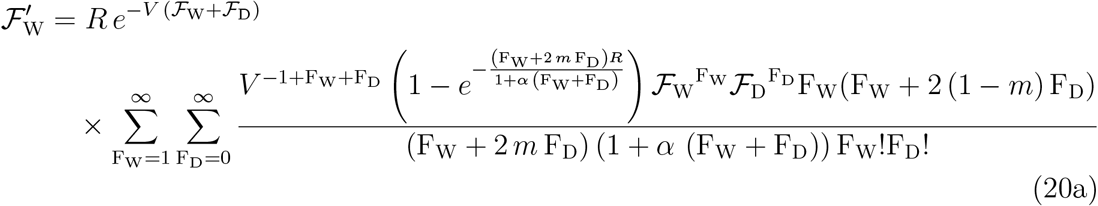

Similar analysis gives 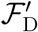, the density of D-mated females in the cloud:

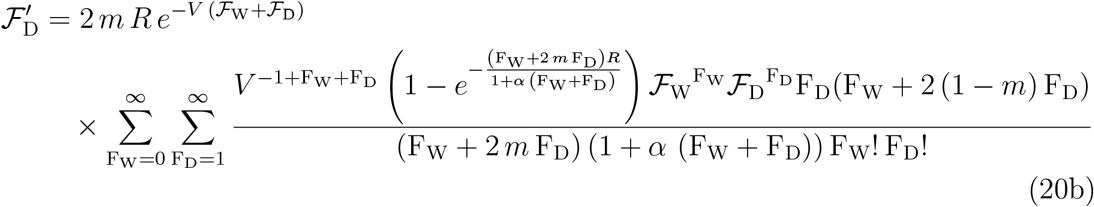

The change of variables 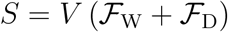 and 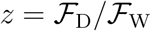 gives:

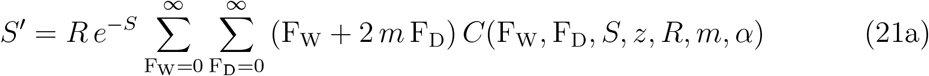

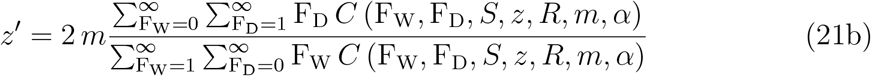

where

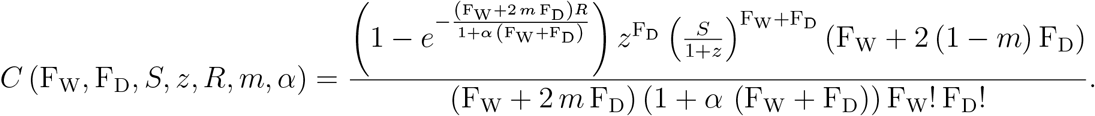

The double infinite sums in (21a) and (21b) can be calculated numerically in an efficient way by noting that the maximum value of the summands occurs at 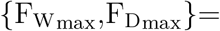 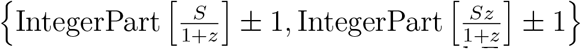 and they decay rapidly for values of F_W_ and F_D_ below and above 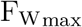 and 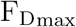, respectively, with the infinite sums thus converging quickly without needing to calculate a prohibitively large number of the coefficients *C*.

In all the scenarios analysed thus far we have seen that the Driving Y tends to fixation and therefore, if *m* is sufficiently high, the population is eliminated. Numerical analysis of (21a) and (21b) shows that is not the case for this scenario. Instead, there is a range of possible outcomes. For most combinations of *R* and *α*, the Driving Y will invade and establish in a population, and then there are three possible outcomes, according to the strength of drive. If *m* is low, then the Driving Y will go to fixation and suppress (but not eliminate) the population. If *m* is somewhat higher, then Driving Y does not eliminate the Wildtype Y, but instead goes to a stable intermediate equilibrium frequency; again, the population is suppressed but not eliminated. Finally, for some values of *R* and *α*, if *m* is higher still, then the frequency of the Driving Y and the population size tend to a limit cycle, oscillating forever. These different behaviours are illustrated in Figure 2, and Figure 3a shows, for a specific value of *m*(= 0.95) the dynamics for different values of *R* and *α*.

**Figure 2:**
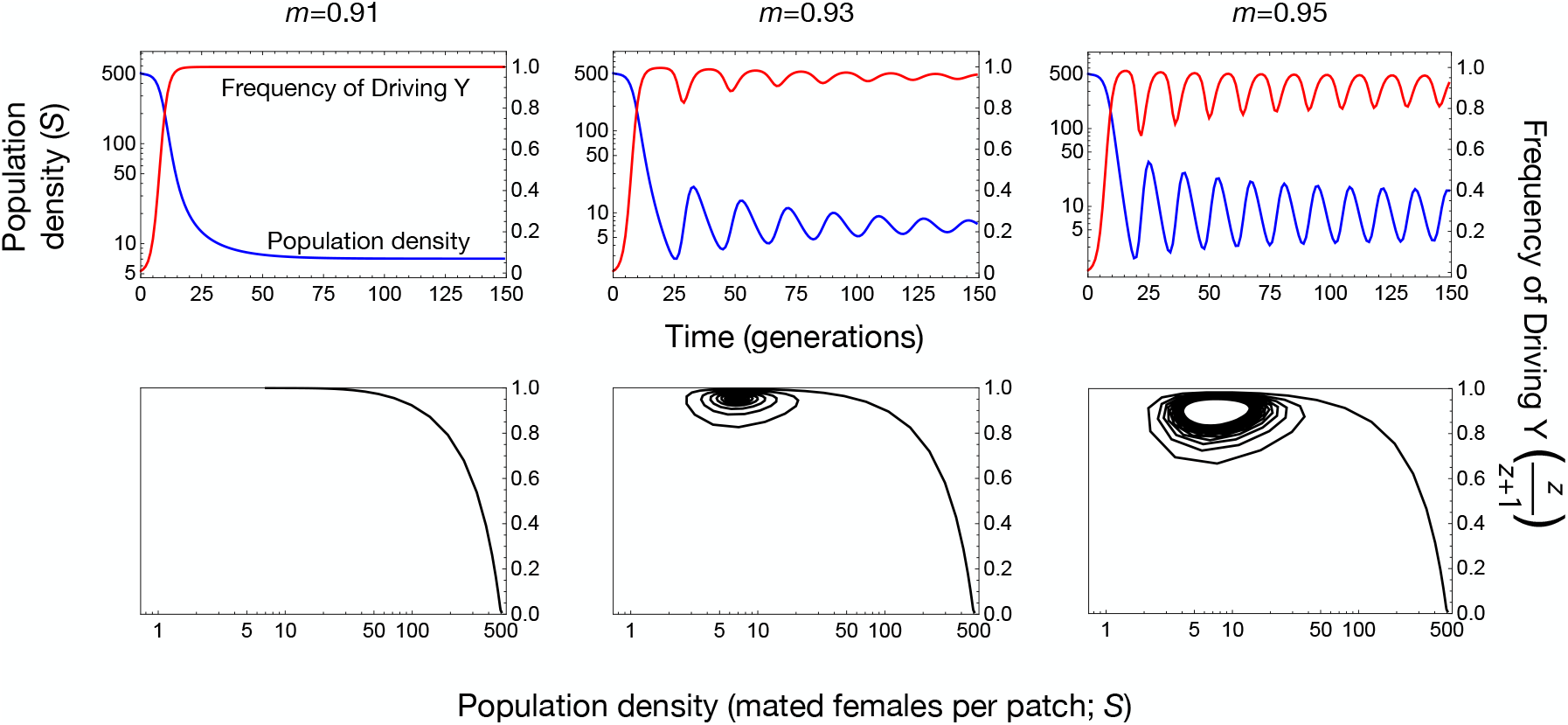
Dynamics and impact of a Driving Y chromosome in a spatial model with local reproduction followed by local mating (Scenario 5). Example dynamics for 3 different strengths of drive (m), shown as time-courses (top; population density in blue, proportion females mated to Driving Y males in red) and as phase planes (bottom), illustrating fixation of the Driving Y (left), an oscillatory approach to a stable intermediate fixed point (middle), and an approach to a stable limit cycle (right). In each case *R* = 6, *α* = 0.01.

**Figure 3:**
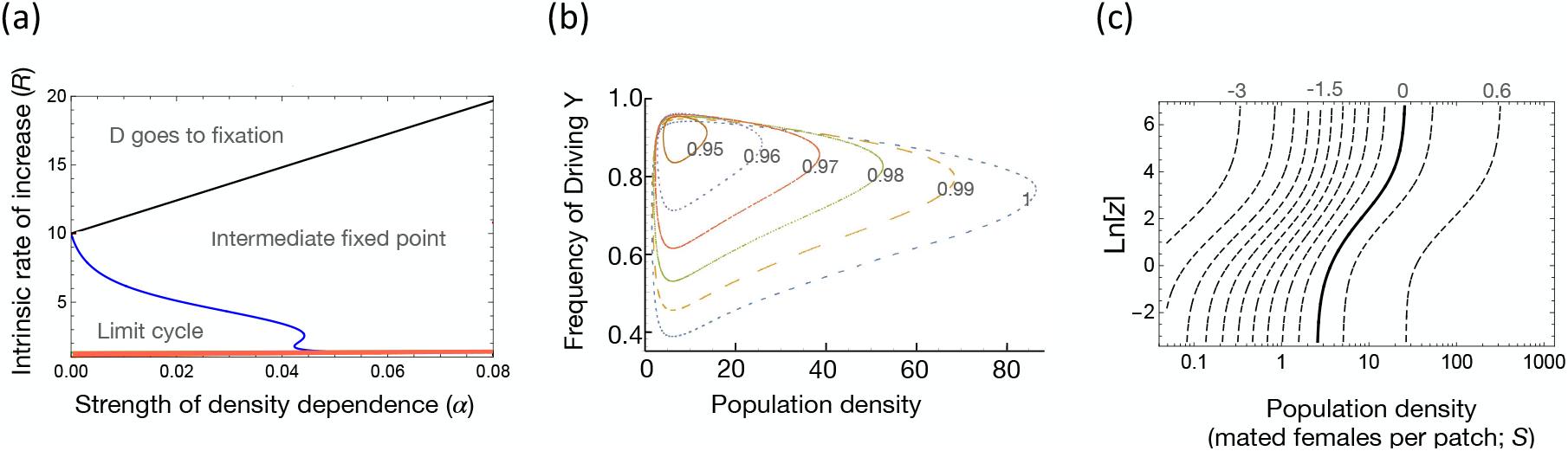
Numerical analysis of Driving Y dynamics with local reproduction followed by local mating (Scenario 5). (a) Critical values of *R* and *α* where the dynamics change from the Driving Y going to fixation, to a stable intermediate fixed point, and to a stable limit cycle, for *m* = 0.95. The red shaded area at the bottom shows the region where the wildtype population exhibits an invasion threshold (i.e., Allee effect), and the effect of a Driving Y can depend on the initial conditions, potentially including population elimination due to the population being driven below its invasion threshold. (b) Closed invariant curves for *m* = 0.95 to 1.00; in each case *R* = 6, *α* = 0.01. (c) Contour plot showing the change across generations in the log-odds that a female has mated a Driving Y male (calculated as Ln[*z*′*/z*], where *z* is the ratio of D- to W-mated females) as a function of the initial log-odds and population density, for *R* = 6, *α* = 0.01 and *m* = 0.95. Solid line shows the 0 contour (no change); contours to the left show negative values (reductions in the D/W ratio), while contours to the right show positive values. Note that all the contours are less than Ln[2 *m*] ≈ 0.642, which is the change in the D/W ratio due to drive, indicating that throughout the investigated parameter space the effect of mating selection is to reduce this ratio.

There is also a region of parameter space in which *R* is sufficiently low that the wildtype population is not able to establish from rare, but instead there needs to be a critical density of females for the population to establish (i.e., it shows a strong Allee effect; Courchamp *et al.* (2008)). Within this region, it seems a population can be eliminated if a Driving Y suppresses it below its invasion threshold, though we have not investigated this phenomenon in detail as it occurs only with relatively small *R*.

For parameter values in which the Driving Y is neither fixed nor lost, linear stability analysis of the fixed point corroborates the simulations. We set *S* = *S*_∞_, z = *z*_∞_ in (21a) and (21b), solve (numerically) for the fixed point {*S*_∞_, *z*_∞_} that corresponds to a given parameter set {*R, m, α*} and then evaluate the 2×2 Jacobian matrix *J* (*S*, *z*; *R, m, α*) of the RHS of (21a) and (21b) at the fixed point to obtain *J*\* = *J* (*S*_∞_, *z*_∞_; *R, m, α*). For an extensive range of parameters {*R, m, α*}, the matrix *J*\* has a conjugate pair of complex eigenvalues λ which indicates the presence of oscillatory dynamics around the fixed point {*S*_∞_, *z*_∞_}. When the modulus |λ| < 1, the fixed point is linearly stable and the variables exhibit dampened oscillations and asymptotically converge to it. When |λ| > 1, the variables oscillate on a unique and stable closed invariant curve that bifurcates from the (unstable) fixed point. The interface between these two regions, i.e., where |λ| = 1, represents surfaces of Neimark-Sacker bifurcation points in the three-dimensional parameter space {*R, m, α*}. We have also shown numerically that the various nondegeneracy conditions associated with Neimark-Sacker bifurcations hold (Kuznetsov, 2004; Khan, 2016) and that the Neimark-Sacker bifurcation is supercritical. For example, one such Neimark-Sacker bifurcation triplet is *{R, m, α}* = {6.0, 0.01, 0.946389}; keeping *R* amd *α* constant, we have |λ| > 1 for *m* > 0.946389 and a unique closed invariant curve exists for every value of 1 ≥ *m* > 0.946389. The size/area of the closed invariant curve (and correspondingly the amplitude of the oscillations in the state variables) increases monotonically from zero at *m* = 0.946389 to a maximum at *m* = 1 (Fig. 3b). The period of the oscillations in the vicinity of the bifurcation point, i.e. at *m* ≃ 0.946389, is ≃2*π*/Im[λ_*m*=0.946389_] = 14.03 and decreases monotonically to 10.30 generations as the amplitude increases as *m* increases from *m* = 0.946389 to 1. There is also a range of parameters {*R, m, α*} (generally for lower values of *m*) where the eigenvalues are real and with modulus < 1; the variables decay monotonically to a stable fixed point and Driving Y fixation.

Further insight can be gotten by considering the ratio 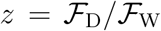, and how that changes from one generation to the next (i.e., *z*′*/z*). In all previously considered scenarios this ratio is equal to 2 *m*, but here it is more complex, and is a function of both the frequency of the Driving Y and, notably, population density, with low densities associated with reductions in the frequency of the Driving Y (Fig. 3c). This total change in the frequency of the Driving Y across a generation can be partitioned between the two relevant events in the life cycle, mating and reproduction. The change in *z* due to reproduction (which isolates the effect of drive) can be quantified by comparing 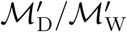, i.e., the ratio of D-males to W-males, to 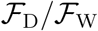, i.e. the ratio of the mated females that gave rise to them. Equations (12) and (13) from Scenario 3 hold here too and show that the quantity 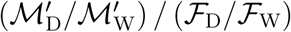 is always 2 *m*, regardless of frequency or density. The change in *z* due to mating is derived from the D/W ratio in mated females to that in the males they had the opportunity to mate with (i.e., 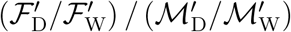, all in the same generation), which isolates the effect of differential mating success or sexual selection. In all cases investigated this ratio is less than 1, indicating a reduction in the D/W ratio, and particularly so at low densities. This is because D-males typically have fewer females to mate with than W-males, because they have fewer sisters, and the difference is greatest at low densities, when sisters are a greater proportion of the potential mates. At the low-density limit, where patches receive at most one female and mating in the next generation will necessarily be between siblings, patches settled by W-mated females will produce more daughters, and therefore more W-mated females, than patches settled by D-mated females will produce D-mated females, and so the frequency of W increases. By contrast, when population density is high and many females settle in each patch, the difference in mating success is much reduced, and the advantage of the Driving Y due to its biased inheritance predominates.

In summary, the one life history scenario we have analyzed in which reductions in density lead to an increased probability of inbreeding shows population persistence regardless of how strong the drive is. This is because Driving Y males have reduced mating success, particularly at low densities, because they have fewer sisters. To further test the hypothesis that it is inbreeding which is protecting the population from elimination, we considered two additional scenarios, in which there is an additional stage of either males or females dispersing before mating (Scenarios 6 and 7, Fig. 1). In either case sib-mating is prevented, and the result, as expected, is the Driving Y goes to fixation, and, for sufficiently high *m*, the population is eliminated (Supplementary Information). Finally, patches in Scenario 5 are arenas for both local density-dependent reproduction and local mating, but it can be shown that only the latter role is needed for the persistence of the Wildtype Y and the population: if reproduction depends on global rather than local density (as if females competed in the cloud for resources that determined their fecundity after settling in patches), the same qualitative outcomes are obtained.

## 3 Homing

To investigate the generality of these results we now consider the same life history scenario (local reproduction followed by local mating, then dispersal) and a completely different form of population suppression gene drive that is autosomal, is transmitted to 1/2 < *d* ≤ 1 of progeny of both male and female heterozygotes, has no effect on the fitness of heterozygotes, and causes homozygotes to die as embryos. Such a gene drive has no effect on the sex ratio, and in non-spatial models (with *d* < 1) it does not tend to fixation in a population, but instead to an intermediate equilibrium frequency, but still can impose a sufficient load on a population to eliminate it.

We consider two types of alleles, the wild type allele W and the drive allele D, that are found in 3 female genotypes, FWW, FWD, FDD, and 3 male genotypes, MWW, MWD, MDD. In this model we assume that both FDD and MDD die as embryos so the only mated females that are possible are WW/WW, WW/WD, WD/WW and WD/WD (using the notation female genotype / male genotype). WW/WW females only produce WW offspring, WW/WD and WD/WW females give WW and WD offspring with proportions (1 – *d*): *d*, and WD/WD females give WW, WD and DD offspring with proportions (1 – *d*)^2^: 2 *d* (1 – *d*): *d*^2^ and, as noted, the DD offspring die early. In this model there is no sex bias so male and female offspring are produced, on average, in equal numbers.

Because transmission rates are equal in the two sexes and there are no heterozygous fitness effects, WW/WD and WD/WW mated females behave identically and can be grouped together so we define 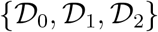 as the cloud densities of WW/WW,(WW/WD + WD/WW) and WD/WD mated females in generation *t*. The number of WW/WW, (WW/WD + WD/WW) and WD/WD mated females settling in a patch is Poisson-distributed with means 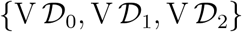, and therefore the probability of having {*D*_0_, *D*_1_, *D*_2_} mated females in a patch is

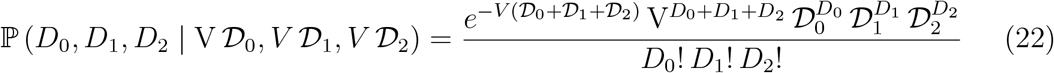

The probability ℙ (*i*_FWW_, *j*_F_ – *i*_FWW_, *i*_MWW_, *j* – *i*_MWW_ – *j*_*F*_ | *j*, *D*_0_, *D*_1_, *D*_2_) that the {*D*_0_, *D*_1_, *D*_2_} mated females generate {*j*_*F*_, *j* – *j*_*F*_} female and male offspring, split as {*i*_FWW_, *j*_*F*_ – *i*_FWW_, *i*_MWW_, *j* – *i*_MWW_ – *j*_*F*_}, is:

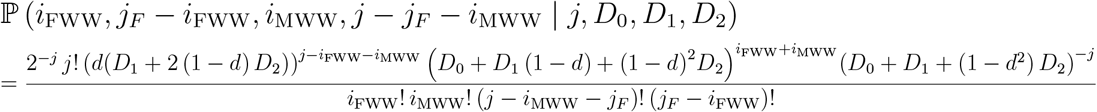

where {*i*_FWW_, *j*_*F*_ – *i*_FWW_, *i*_MWW_, *j* – *j*_*F*_ – *i*_MWW_} are the numbers of FWW, FWD, MWW and MWD (viable) offspring, respectively (FDD and MDD offspring die as embryos and we only focus on viable offspring).

Female and male offspring in the patch are paired randomly to generate mated females of WW/WW, (WW/WD + WD/WW) and WD/WD types. By averaging over all possible probability-weighted values of *i*_FWW_, *i*_MWW_ and *j*_*F*_, we obtain the expected numbers of mated females in the patches, conditional on {*j*, *D*_0_, *D*_1_, *D*_2_}:

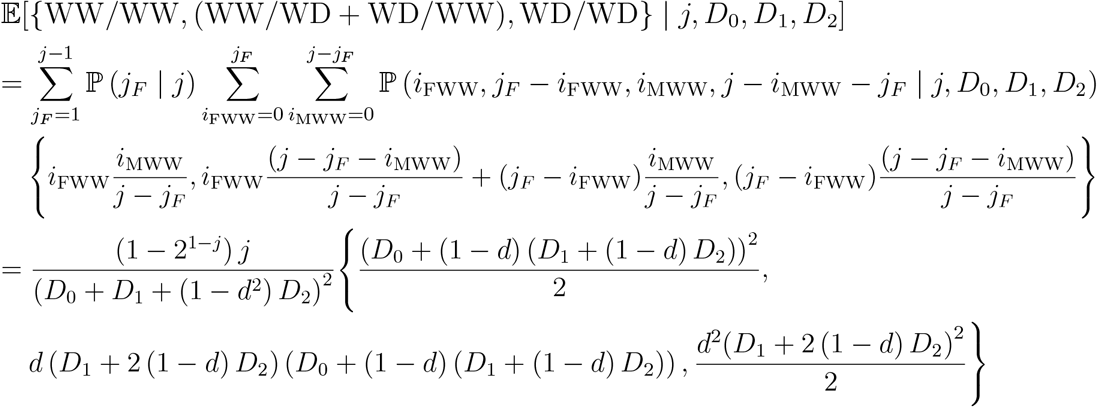

where 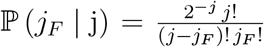 is the probability of having *j*_*F*_ female offspring out of *j* total offspring.

In this model all types of mated females generate offspring, but the *D*_2_ WD/WD females in a patch only generate, on average, a fraction of (1 – *d*^2^) viable offspring (the remainder of the offspring, namely FDD and MDD, die as embryos and thus do not compete with other genotypes). The total number of offspring produced in a patch is Poisson-distributed and the probability of having generated *j* viable offspring in total in a given patch is then

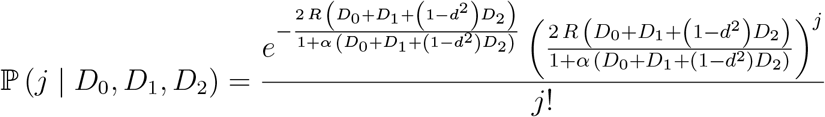

and the expected numbers of mated females in the patches, conditional on {*D*_0_, *D*_1_, *D*_2_}, is obtained by averaging over all probability-weighted values of *j* from 2 to infinity:

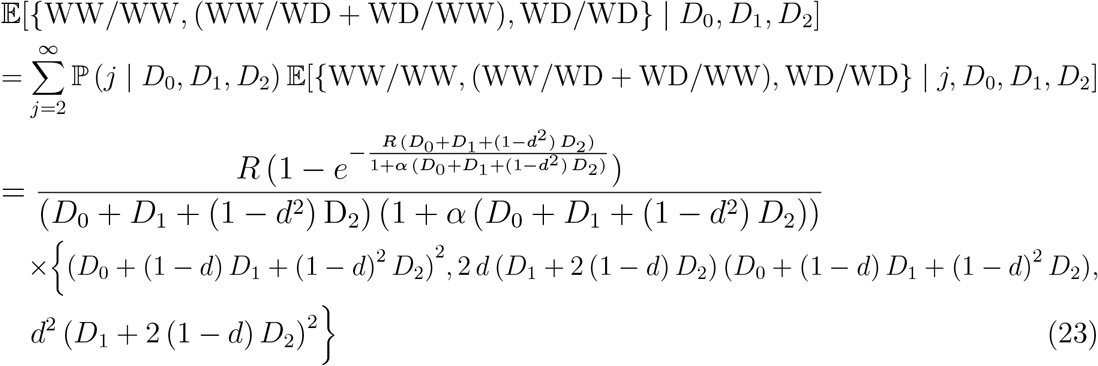

At this stage the mated females in each patch migrate to the cloud. In order to calculate the densities 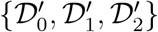 of the newly mated females in the cloud for generation *t* + 1, we average the expected numbers of newly mated females in a patch, calculated in (23), over all values {*D*_0_, *D*_1_, *D*_2_} of mated females that arrived in the patches from the cloud during generation *t*, weighted by 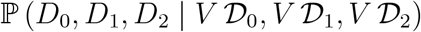 in (22).

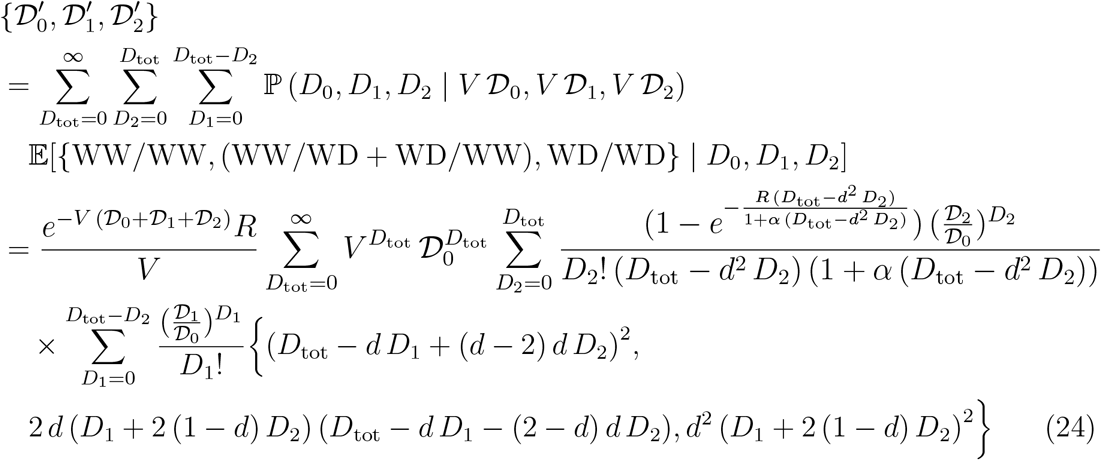

where *D*_tot_ = *D*_0_ + *D*_1_ + *D*_2_.

The innermost sum in *D*_1_ can be calculated analytically so (24) for 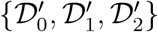 reduces to an outer infinite sum in *D*_tot_ and an inner finite sum in *D*_2_:

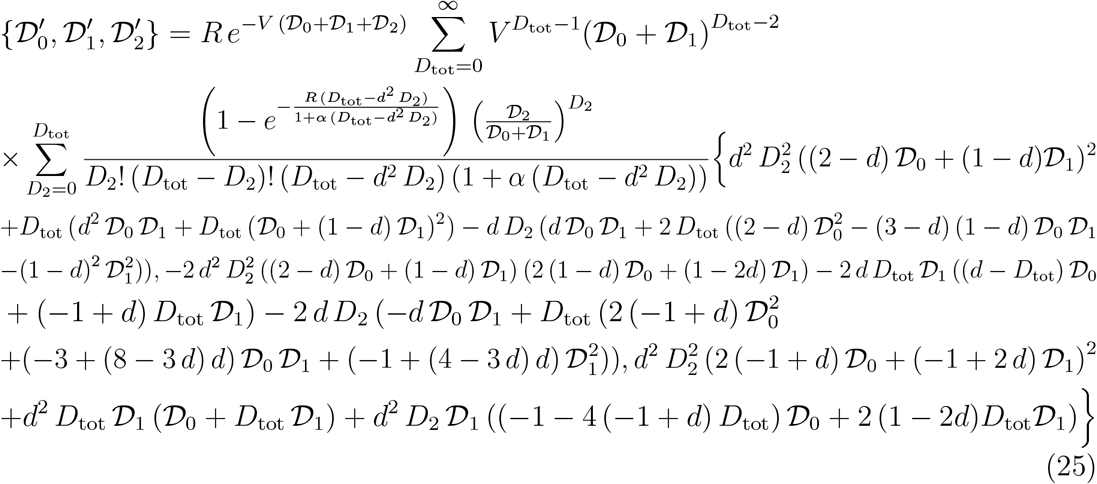

The recurrence vector equation (25) is sufficient to describe the dynamics of this system and is used to calculate the densities of mated females in the cloud from one generation to the next.

To aid understanding, we present the results of the model in terms of 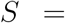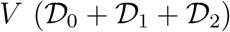, the (global) average number of mated females per patch, 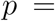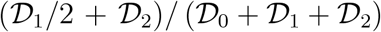, the frequency of D heterozygotes that participated in the matings (and twice the frequency of the D allele itself), and 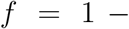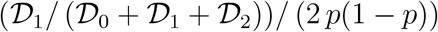, analogous to the standard inbreeding coefficient, except it measures the correlation of mates rather than of fusing gametes. If we iterate the transition equations using different parameter values and initial conditions then, assuming the pure wildtype population does not have an invasion threshold, the driver typically either goes to a stable fixed point or, for stronger drive, to a stable limit cycle, and in either case the population persists, regardless of how strong the drive is. Example dynamics are shown in Fig. 4a.

**Figure 4:**
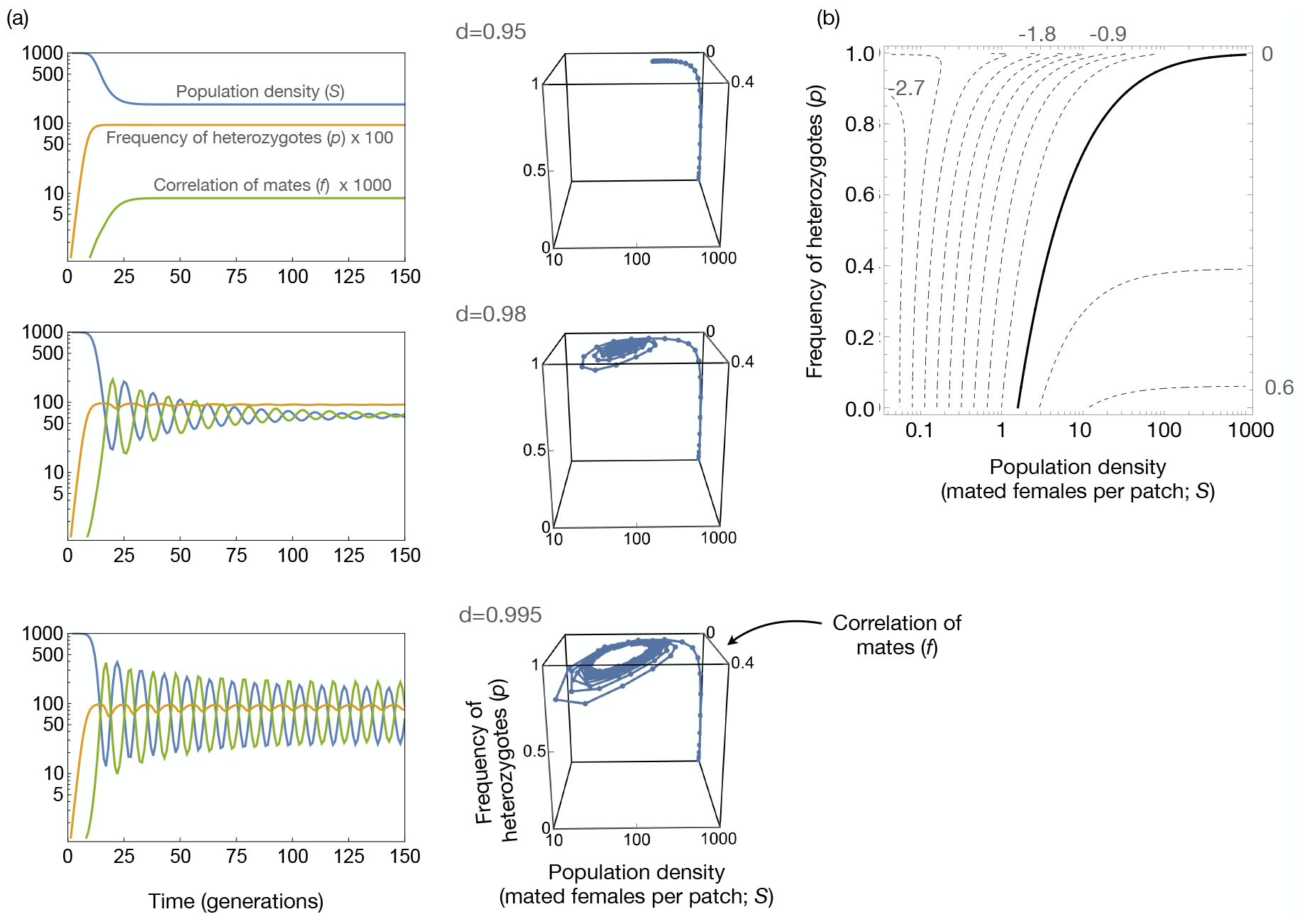
Population persistence in spatial models of a homing construct with local reproduction followed by local mating (same as in Scenario 5 for the Driving Y). (a) Example time courses and phase plots for *R* = 6, *α* = 0.005 and three different strengths of drive *d*. For ease of viewing the time courses, the heterozygote frequency has been multiplied by 100 and the correlation of mates by 1000. (b) Contour plot showing how the natural log of the ratio of construct frequency in generation *t* + 2 to that of generation *t* + 1 depends on the population size and frequency of heterozygotes in generation *t*. Changes calculated for an initial correlation of mates of *f* = 0.2; other values of *f* give comparable results. Calculations are for *d* = 0.995, *R* = 6, *α* = 0.005.

Again, we can partition the total change in frequency of the construct due to the various processes occurring through the life cycle. In this case, it is more convenient to measure changes in raw frequencies. There is no differential mating success, so the two relevant processes are drive and differential survival due to the death of DD embryos. It can be shown that if *p* and *f* are defined for the adults of one generation, then the ratio of the frequency of the D allele in the zygotes they produced to that in the adults (*p*), which isolates the effect of drive, is 2 *d*, and the ratio of the frequency in the next generation of adults to the zygotes from which they were derived, which isolates the effect of differential mortality, is (1 – *dp* – *d* (1 – *p*) *f*)/(1 – *d*^2^ *p* (*p* + (1 – *p*) *f*)). As expected, it is always <1.

Note that population size does not have an immediate impact on the change in construct frequency due to drive or mortality selection (it does not appear in the above expressions), but it does have a delayed effect. In particular, a smaller population density in generation *t* leads to a larger correlation between mates (*f*) in generation *t* + 1, because there is an increased frequency of mating between siblings. This larger correlation in generation *t* + 1 leads to a larger reduction in the frequency of the driver from generation *t* + 1 to generation *t* + 2, because of the lower productivity of WD/WD mated females, which in turn is due to the differential embryonic mortality – the death of DD embryos – in generation *t* + 2. This delayed inverse density-dependent selection against the driver is illustrated in Fig 4b.

Thus, though the details differ from the case of Driving Y, the overall result remains the same: an increased frequency of inbreeding at low population densities leads to increased selection against the driver, reducing its frequency and allowing the population to persist, regardless of how strong the drive is.

## 4 Inbreeding depression

Our analyses have demonstrated that when reductions in population density lead to an increase in inbreeding, that can increase the natural or sexual selection against the driver and allow the population to persist. However, inbreeding can only rescue a population to the extent that the inbred progeny are themselves fit enough to contribute. Thus far we have assumed no difference in fitness between inbred and outcrossed progeny. To further test the hypothesis that inbreeding plays a central role in the observed dynamics, we now allow for inbreeding depression, in which inbred progeny have reduced fitness. At the limit of inbred progeny being completely inviable or sterile, we might expect the dynamics to revert to population elimination.

### 4.1 Driving Y

This scenario involves the same local mating, global mixing of mated females, and local reproduction steps as Scenario 5. However, in this model some or all of the females mated by sibling males are sterile and are thus removed from the mated females that disperse to new patches for local reproduction. We define the inbreeding depression coefficient 0 ≤ *δ* ≤ 1 as the probability a sibling-mated female is sterile; *δ* also represents the fraction of sibling-mated females that is removed from the ensemble of mated females that travel to the cloud and then disperse into patches (mathematically, there is no difference between removing the sterile mated females before or after they travel to the cloud). As in Scenario 5, the F_W_ + F_D_ mated females generate *j* offspring in total in the patch. However in this scenario, the *j* offspring are made up of 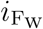 females from W-mated mothers, 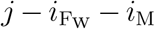 females from D-mated mothers, *i*_2_ W-males and *i*_M_ – *i*_2_ D-males (where *i*_M_ = *i*_2_ + *i*_3_ is the total number of male offspring). The probability of 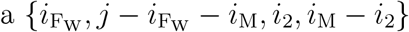 quadruplet is derived from a multinomial distribution with *j* trials and normalised weights 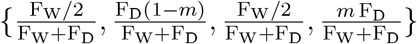, since, on average, fractions of 1/2 and 1 – *m* of W-mated and D-mated females’ offspring are female, with the rest of the offspring being W- and D-males, respectively.

Hence, the probability 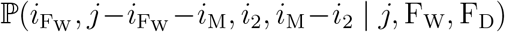 of having 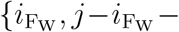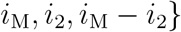 offspring in the patch (conditional on *j* total offspring from {F_W_, F_D_} mated females) using the weights above is

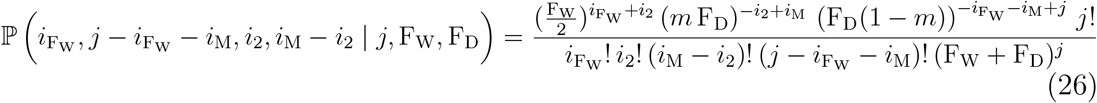

In order to remove, on average, a fraction 0 ≤ *δ* ≤ 1 of sibling-mated females from the total number of new mated females in the patch, we present a proof by induction in the Supplement (Section 7.3) that the expected number of new sibling-mated W-mated females in the patch, conditional on 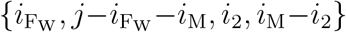 offspring from {F_W_, F_D_} mated females, is 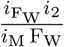. A similar proof, not shown, gives the expected number of new sibling-mated D-mated females in a patch, conditional on 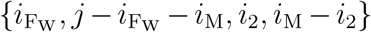 offspring from {F_W_, F_D_} mated females, as 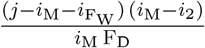.

As a result, the expected number of new fertile W-mated females in a patch, conditional on 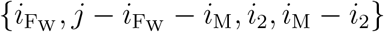 offspring from {F_W_, F_D_} mated females is:

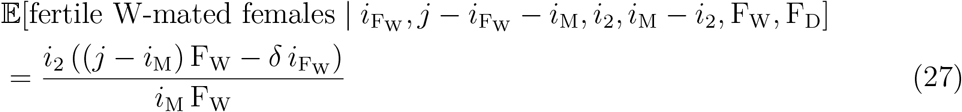

(the probability that each new sibling-mated female is sterile is *δ*, so we removed a fraction *δ* of the sibling-mated females’ contribution to the total number of expected number of fertile mated females in the patch).

To obtain the expected number of W-mated females in the patch, conditional on {*j*, F_W_, F_D_}, we now sum over all possible 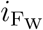, *i*_M_ and *i*_2_, noting that *i*_M_ ≥ *i*_2_ ≥ 1 to ensure the presence of at least one W-male and *j* – 1 ≥ *i*_M_ to ensure the presence of at least one female:

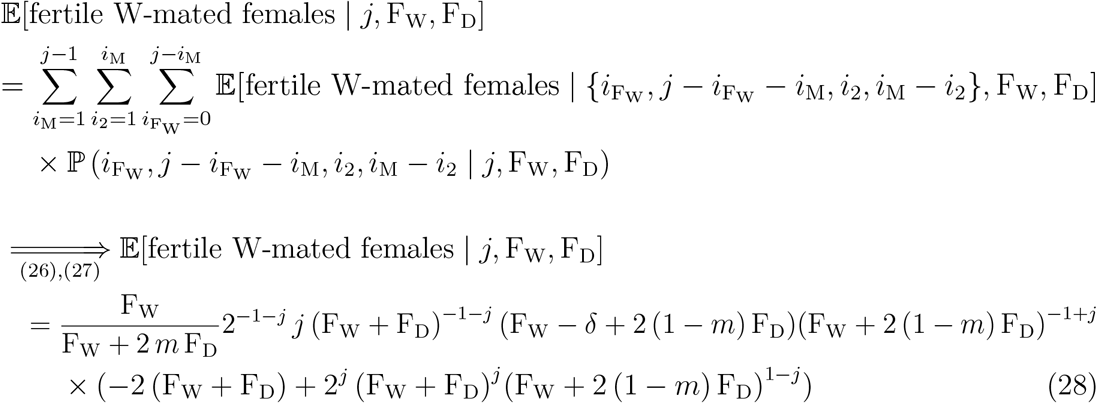

At this stage the fertile mated females in each patch migrate to the cloud. In order to calculate the density of W-mated females in the cloud, we combine (7) for the probabilities 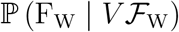 and 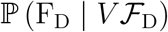 of having F_W_ and F_D_ mated females in a patch, (8) for the probability ℙ (*j* | F_W_, F_D_) that the F_W_ + F_D_ mated females generate *j* offspring in total in the patch and (28) for the expected number of W-mated females, conditional on *j* offspring from F_W_ and F_D_ mated females in the patch. Their product is then summed over all possible values of F_W_, F_D_ and *j* to give the average number of W-mated females across all patches, and is then divided by *V*, to give the expression for 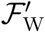, the density of W-mated females in the cloud in generation *t* + 1 (the innermost summation over *j* is calculated analytically so the result below is in terms of a double infinite sum over F_W_ and F_D_):

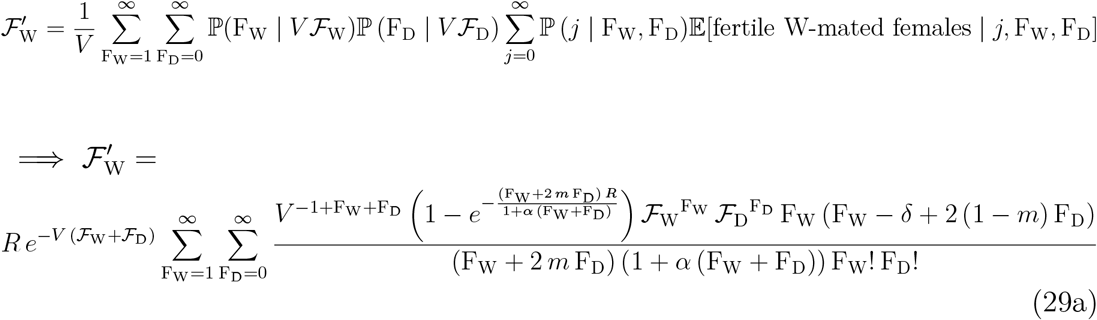

Similar analysis gives 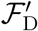, the density of D-mated females in the cloud:

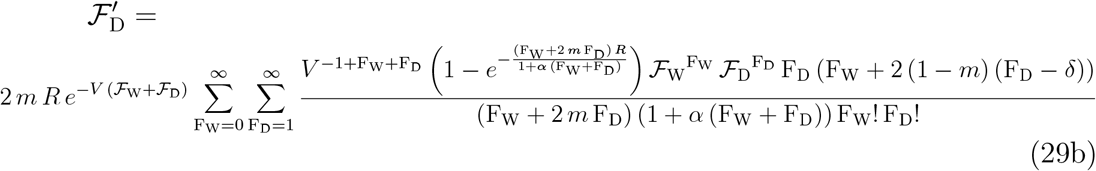

The RHSs of (29a) and (29b) contain a negative term, proportional to *δ*, which represents the population suppression effect due to inbreeding. When *δ* = 0, i.e. when the sibling-mated females are all fertile, (29a) and (29b) reduce to (20a) and (20b).

We focus on the extreme case of *δ* = 1 (i.e., females mated to their brothers produce no viable offspring) and introduce the change of variables 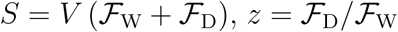 and F_W_ = F – F_D_. We will show that the drive always goes to fixation by showing that the quantity 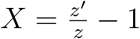, derived from the equations above, is always positive:

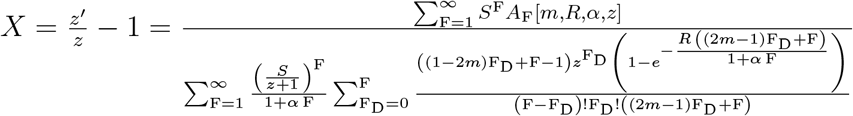

where the F-th coefficient *A*_F_ = *A*_F_ [*m, R, α, z*] is

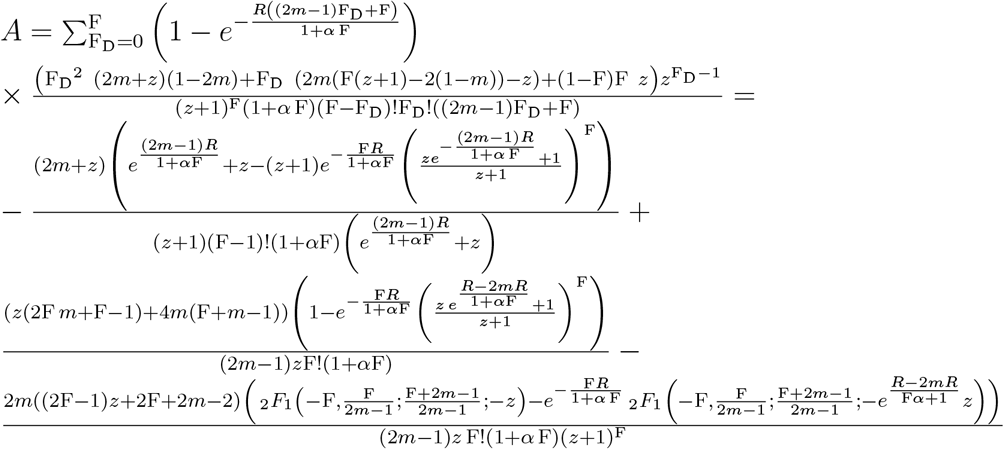

where _2_*F*_1_(−F, *a*; *a* + 1; *x*), F ϵ ℕ is a polynomial of order F in *x* and is a special case of the Gauss Hypergeometric Function.

The denominator of *X* above is the transformed expression for 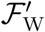, so it is always positive and can be ignored. We postulate that all the coefficients *A*_F_ in the numerator are positive so that *X* is always positive. For a given set of parameters *m*, *R*, *α*, *A* only depends on *z* ϵ [0, ∞), i.e. it does not depend on *S*. At *z* = 0,

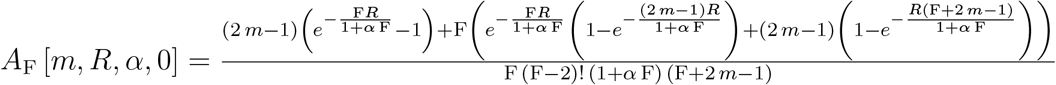

which is clearly positive for all *m*, *R*, and *α* It can also be shown analytically that *A*_F_ [*m*, *R*, *α*, *z* → ∞] → 0 for all *m*, *R*, and *α*. We have evaluated *A*_F_ [*m*, *R*, *α*, *z*] for a large set of *m*, *R*, and *α* and we always find that it decreases monotonically from *A*_F_ [*m*, *R*, *α*, *z* = 0] to 0 asymptotically as z → ∞. Based on our analysis, we postulate that *A*_F_ [*m*, *R*, *α*, *z*] > 0 for all values of *m*, *R*, *α* and *z*. If so, it also means that 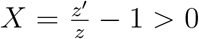 for all values of *m*, *R*, *α*, i.e. the ratio 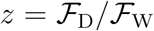 is always greater in the next generation, irrespectively of the values of *S* and *z* (or 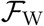 and 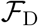) in the current generation. This can only result in fixation of the Driving Y as *t* → ∞, and, if *m* is sufficiently large, population elimination (Fig. 5a). Numerical analysis suggests that *z* does not go to infinity for *δ* < 1, but still there is a large parameter range in which the population is eliminated.

**Figure 5:**
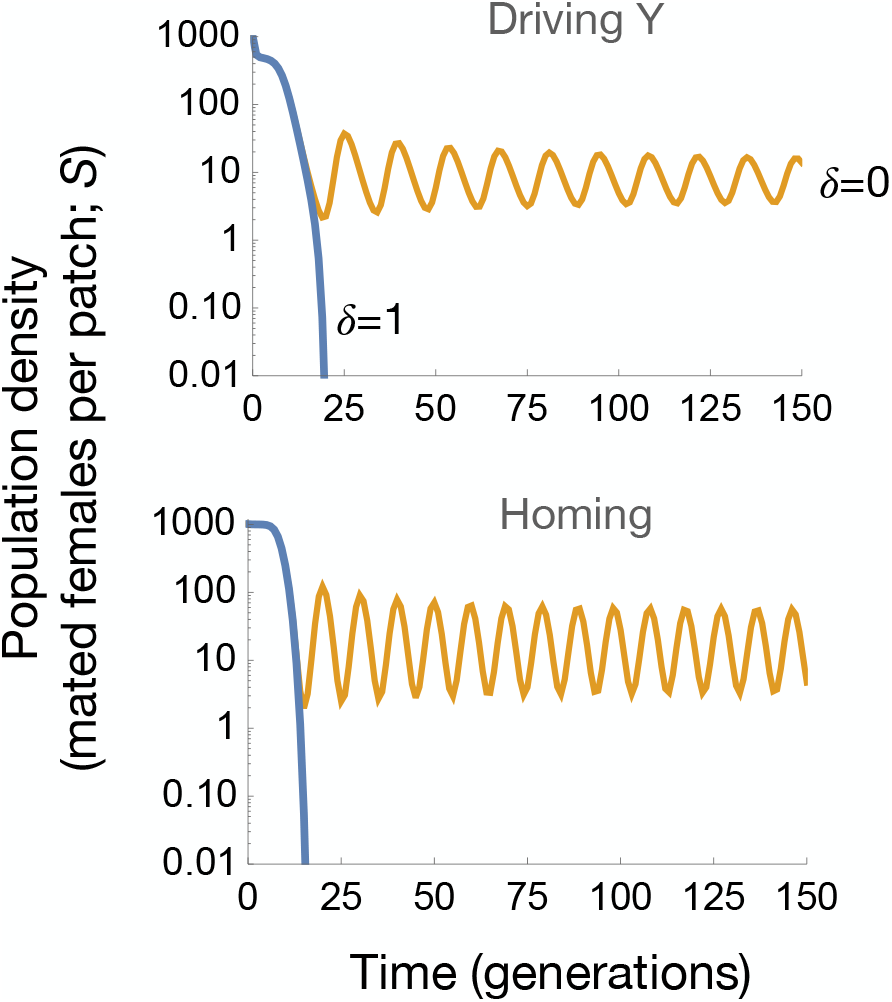
Time courses for population density after the release of a gene drive, demonstrating that strong inbreeding depression (*δ* = 1) can lead to population elimination. Top: A Driving Y (*R* = 6, *α* = 0.01, *m* = 0.95). Bottom: A homing construct (*R* = 6, *α* = 0.005, *d* = 1).

### 4.2 Homing

We now investigate the effect of inbreeding depression on the dynamics of a homing construct. For the sake of simplicity, we will focus on the case of *d* = 1, i.e. where all the progeny of WW/WD and WD/WW mated females are WD males and females and all the progeny of WD/WD mated females are DD males and females, while, as before, all the progeny of WW/WW mated females are WW males and females. As we assume that DD males and females die as embryos, in this limit case of *d* = 1 we can ignore WD/WD mated females as they do not produce viable offspring.

We therefore consider 3 types of mated females: non-sibling-mated WW/WW females, sibling-mated WW/WW females and (WW/WD + WD/WW) mated females. The latter are all non-sibling-mated as they represent pairings between a WW male or female (i.e. an offspring of WW/WW mother) and a WD female or male (i.e. an offspring of a (WW/WD + WD/WW) mother), which cannot ever be siblings as they are produced from two different types of mother.

We define as 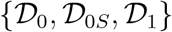 the cloud densities of non-sibling-mated WW/WW, sibling-mated WW/WW, and (WW/WD + WD/WW) mated females in generation *t*. Any volume *V* in the cloud contains *on average* 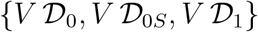 numbers of individuals. In this model, the mated females in the cloud settle into patches with *average* numbers of individuals equal to 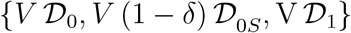. The cloud density 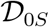 of sibling-mated females is derived from the entirety of the sibling-mated females in the patches. We then only allow a fraction 1 – *δ* of these females to disperse into patches.

We have assumed that the sibling-mated females that settle in the patches are the (fully) fertile portion of the sibling-mated WW/WW females in the cloud, so once settled in patches, they are indistinguishable from the non-sibling-mated WW/WW females. They can therefore be combined inside each patch into a single cohort of WW/WW mated females. A random sample of {*D*_0_, *D*_1_} WW/WW and (WW/WD + WD/WW) mated females, different for each patch and Poisson-distributed with means 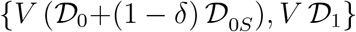, is drawn from the cloud and settles in each patch (the mean number of WW/WW females that settle in a patch is the sum of non-sibling mated and *fertile* sibling mated females in the cloud). The probability of having {*D*_0_, *D*_1_} mated females settle in a patch is

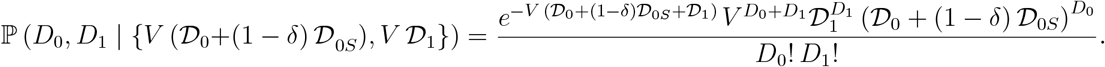

The mated females produce on aggregate *j* offspring. The number of successful offspring that each mated female produces is Poisson-distributed with a mean 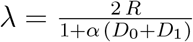. The probability that the *D*_0_ + *D*_1_ mated females generate *j* offspring in total in a given patch is then

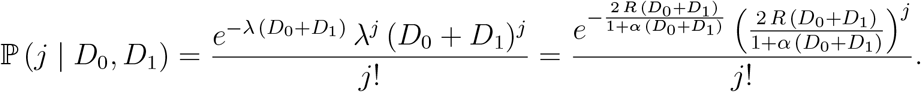

Of the *j* offspring, *j*_*F*_ are female and *j* – *j*_*F*_ are male. The WW/WW females produce {*i*_FWW_, *i*_MWW_} WW female and male offspring and the (WW/WD + WD/WW) females produce {*j*_F_ – *i*_FWW_, *j* – *j*_F_ – *i*_MWW_} WD female and male offspring. The conditional probability of having {{*i*_FWW_, *i*_MWW_}, {*j*_F_ – *i*_FWW_, *j* – *j*_*F*_ – *i*_MWW_}} offspring is:

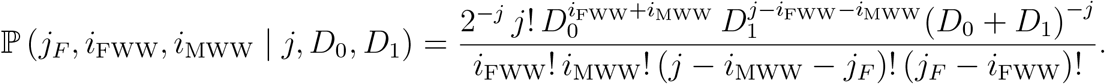

Each female offspring in the patch chooses a random male partner and the resulting mated females fall into 3 categories: non-sibling-mated WW/WW, sibling-mated WW/WW and (non-sibling) (WW/WD + WD/WW) females. As shown in the Supplement for the Driving Y case, the expected fraction of new sibling-mated females from offspring of *n* mothers is 1/*n* of the total new mated females (and, as a result, the fraction of non-sibling mated females is (*n* – 1)*/n*). The *i*_MWW_ WW male and *i*_FWW_ WW female offspring of *D*_0_ WW/WW females in the patch produce, on average, 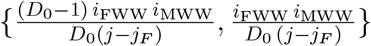 non-sibling- and sibling-mated WW/WW females. The expected number of each category, conditional on {*D*_0_, *D*_1_} mated females settling in the patch, is thus derived by averaging 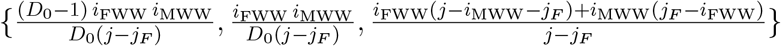 over all the probability weighted values of *j*, *j*_*F*_, *i*_FWW_, and *i*_MWW_:

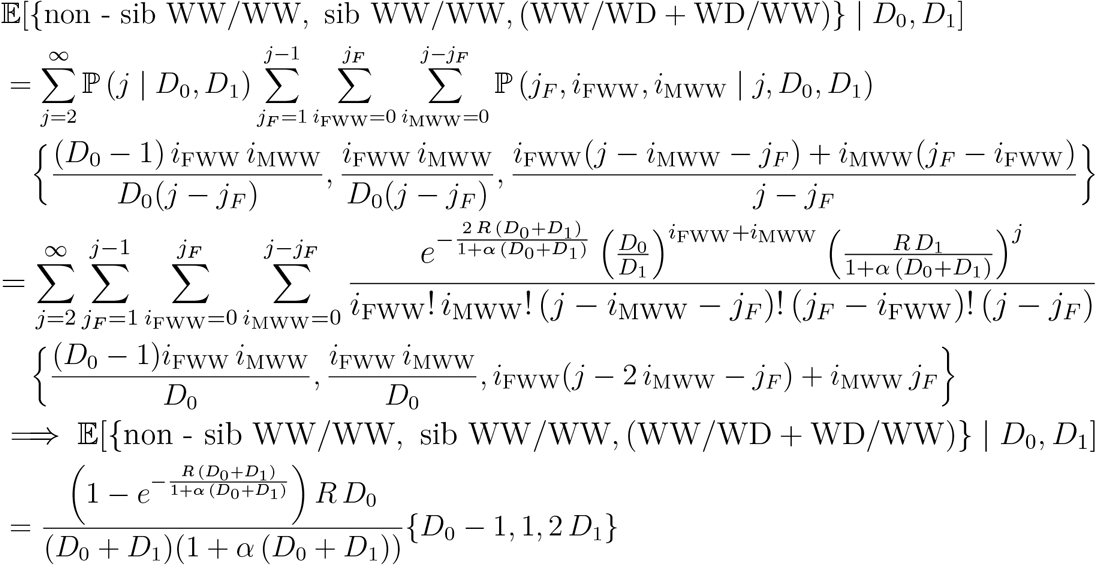

At this stage, the mated females in each patch migrate to the cloud. In order to calculate the densities 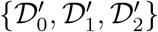 of the newly mated females in the cloud for generation *t* + 1, we average the expected numbers of newly mated females in a patch, calculated above, over all probability-weighted values {*D*_0_, *D*_1_} of mated females that arrived in the patches from the cloud during generation *t* (and divide by V):

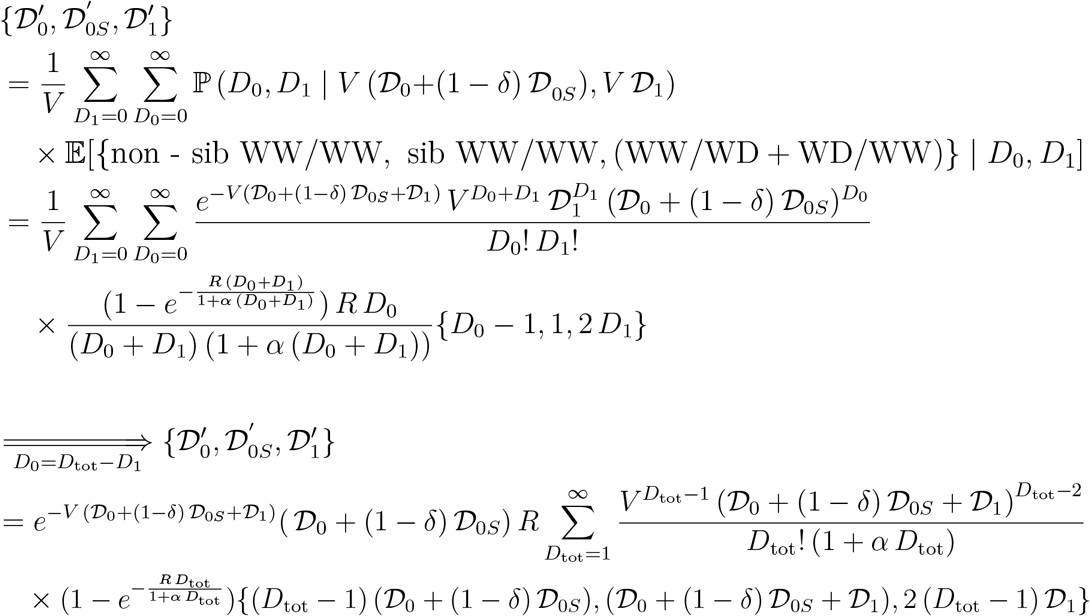

where the variable change *D*_0_ = *D*_tot_ – *D*_1_ turns one of the infinite summations into a finite one which is computed analytically, leaving only a single infinite summation in 1 ≤ *D*_tot_ < ∞.

We now focus on the case of *δ* = 1, i.e. where none of the offspring from sibling matings survive. Sibling-mated females can thus be ignored and the equation above simplifies to

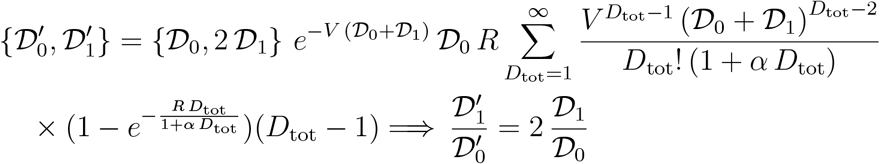

The result above, i.e. the doubling of the ratio 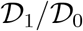 with every generation, means that as *t* → ∞ the cloud will only contain (WW/WD + WD/WW) mated females which in turn only produce non-viable DD offspring (given that *d* = 1). The population will thus asymptotically go to 0 (Fig. 5b).

## 5 Discussion

The key role of sex and breeding system in the strength and consequences of gene drive is well established, having been studied from theoretical, experimental and comparative perspectives (Burt & Trivers, 2006; Agren & Clark, 2018). It is therefore reasonable to expect that if a driver suppresses a population and that leads to an increased frequency of inbreeding, then there may be a limit to how far the suppression can go, regardless the strength of the drive (Bull *et al.*, 2019). Here we have presented modelling in support of this reasoning.

First, we considered the fate of a Driving Y under different life history scenarios. In a non-spatial model in which both mating and reproduction occur in a well-mixed cloud (scenario 1), a Driving Y will gradually replace the Wildtype Y and go to fixation, and, if drive (*m*) is high enough (relative to the population growth rate *R*), the population will be eliminated. If reproduction occurs in the cloud and mating in patches (scenario 2), or vice versa (scenario 3), or if individuals mate and reproduce in a patch followed by the offspring dispersing (scenario 4), then there is no qualitative difference in the dynamics: the Driving Y goes to fixation and, if *m* is sufficiently high, the population is eliminated. However, if the order of activities within patches is reversed, so that mated females settle in patches and reproduce and then the offspring mate before returning to the cloud (scenario 5), then there is a qualitative difference: the Driving Y will only fix for *m* below a threshold value, and otherwise remains polymorphic, and the population is suppressed but not eliminated regardless of how high *m* is. Conversely, if this scenario is modified by interposing another round of dispersal of either males or females between reproduction and mating (scenarios 6 and 7, see SI), then again the Driving Y goes to fixation and, if *m* is sufficiently high, the population is eliminated. Thus, the only life history scenario in which the probability of inbreeding increases at low densities is the one at which allows indefinite persistence of both the wild type Y chromosome and the population. This scenario has the same life cycle as Hamilton’s (1967) much studied local mate competition model of sex ratio evolution, the difference being in the ecology, where he considered the population size (number of females per patch) to be exogenously fixed, we consider it a dynamic variable responding to the presence of the Driving Y.

These results can be interpreted in terms of altruism: a Wildtype Y is altruistic (compared to a Driving Y) in the sense of foregoing transmission to allow the production of more females. That can be a useful thing to do if those extra females mate with the W-males, but otherwise not. When population sizes are large, with many mated females settling in a patch, the extra females produced by a W-male’s forbearance are shared out equally among all the males in the patch, and so the W gains relatively little, not enough to compensate for the reduced transmission. However, if population sizes are very low, with at most 1 mated female settling in a patch, then the extra daughters produced by the W-male all go to his W-bearing sons, and the frequency of W increases.

Second, we have shown that the same life history scenario leads to the same qualitative outcome (population persistence regardless of the strength of drive) for a gene drive construct using the homing reaction to knock out an essential gene, though the precise details differ. For a Driving Y, increased inbreeding means that the number of sisters a male has is an important component of his fitness, and, unavoidably, Driving Y males will have fewer sisters than wildtype males. For autosomal drivers causing recessive lethality, increased inbreeding means wildtype (WW) individuals are mating with wildtypes, and drivers (WD) with drivers. In the absence of inbreeding depression, mating with a sibling is more productive for wildtypes, producing a full complement of offspring, than for drivers, who will be at increased risk of mating with another driver, in which case only a fraction (1 – *d*^2^) of their progeny will be viable.

Third, we have shown for both types of drive that if inbred progeny are prevented from contributing to the population (by imposing strong inbreeding depression), then the previous advantage of the wildtype at low density disappears and the results change again, with sufficiently strong drive once more able to eliminate the population. For a Driving Y, population persistence relies on the wildtypes having an advantage at low densities because they can mate with their sisters, but if those matings do not produce viable offspring, the advantage disappears. Similarly, for a homing construct, the wildtype can have an advantage at low densities because mating between relatives does not carry the risk of producing lethal DD offspring, but they are lethal just by being inbred then again the advantage disappears.

Thus our modelling suggests that populations can persist in the face of strong gene drive, even in the absence of resistance, if three requirements are met: the target population shows spatial structure; reductions in population density lead to an increased probability of inbreeding; and inbred progeny have sufficiently high fitness. The extent to which the three criteria exist in a particular target species will need to be assessed on a case-by-case basis. If the population is not eliminated, then it can still be significantly suppressed, and this may be sufficient by itself for the purposes, or may be a useful component of a multi-pronged elimination programme. In principle, populations may also be rescued by selection for genetic variants that increase the frequency of inbreeding independently of density, though, again, strong inbreeding depression will militate against such an effect (Bull, 2016; Bull *et al.*, 2019).

Inbreeding depression in our model reduces the population growth rate at low densities, and therefore acts as an Allee effect (Luque *et al.*, 2016). Even in our baseline model, without inbreeding depression, there is a small region of low population growth rates where the wildtype population shows a strong Allee effect, requiring a threshold density to establish. This effect arises because at low densities, and low values of *R*, a single female may not produce any sons to mate her daughters. Within this region of parameter space it is possible for a Driving Y to suppress the population below the threshold density and thereby eliminate it. The effect of including inbreeding depression is to increase the region of parameter space in which a wildtype invasion threshold density exists and elimination is possible. Most or all species show inbreeding depression, primarily due to the unmasking of deleterious recessive mutations, and, all else being equal, the magnitude of the effect is expected to be greater in populations that previously were large and outcrossed (Tanaka, 2000; Frankham, 2005; Charlesworth & Willis, 2009). Inbreeding depression is not the only possible source of an Allee effect: for example, low densities can also lead to difficulties in finding a mate (Courchamp *et al.*, 2008). In our model we have assumed that if there is a single male in the patch, then all females will get mated, but for many species this assumption may not be valid. We would expect that Allee effects due to difficulties in finding a mate (or any other source) could also tip the balance from population persistence to elimination (see also Dhole *et al.* (2020)). Interestingly, there will often be synergistic interactions between genetic and ecological Allee effects (Wittmann *et al.*, 2018). The possibility of exploiting Allee effects for pest control more generally has been previously discussed (Liebhold & Bascompte, 2003; Blackwood *et al.*, 2018).

The interaction of gene drive and spatial processes have been modelled in many ways, revealing a diverse array of effects (Dhole *et al.*, 2020). In deterministic partial differential equation models with local diffusion, sufficiently strong drive leads to population elimination, though it takes longer than in a panmictic population (Beaghton *et al.*, 2016). On the other hand, stochastic spatial models have shown that populations can persist even with arbitrarily strong drive, and identified three types of effect protecting the population from elimination. First, in some cases it may be that the connectedness of populations across the landscape is such that a drive, released in one part of the landscape, does not reach some specific refugia populations before it itself goes extinct (North *et al.*, 2013; Eckhoff *et al.*, 2017). This effect can be particularly acute in highly seasonal environments, where a prolonged and severe dry season can lead to (transient) population isolation, and a driver might reach a locale during the wet season, but not attain a sufficiently high frequency to survive through a dramatic dry season bottleneck (Eckhoff *et al.*, 2017; North *et al.*, 2019, 2020). In principle, the issue of refugia can be addressed by more widespread releases, appropriately timed for the beginning of the wet season (Lambert *et al.*, 2018), ensuring the drive is introduced into all parts of the landscape. Second, even if populations are sufficiently connected that the gene drive eventually gets to all parts of the landscape, the population may nonetheless persist because the wildtype is able to colonize previously cleared areas, and grow in abundance, it taking some time for the driver to get there and suppress the population, by which time the wild type has spread to another previously cleared location, resulting in a phenomenon which has variously been referred to as “dynamic metapopulations” (North *et al.*, 2019), “colonization-extinction” dynamics (North *et al.*, 2020), and “chasing” dynamics (Godfray *et al.*, 2017; Champer *et al.*, 2021). Finally, in the model presented in this paper we have seen that even with 100% global dispersal every generation, spatial processes can protect a population from elimination if low densities lead to increased inbreeding and inbred progeny are sufficiently fit, because it leads to selection against the driver.

In each of these three cases there is something that keeps the wildtype and driver alleles from direct maximal competition, be it refugia on an insufficiently connected landscape, or a small spatial separation between colonizing wildtypes and chasing drivers, or the random assortment of females into patches at low densities. We have demonstrated that the last mechanism relies on inbreeding – the ability of brothers and sisters to successfully mate and reproduce – and suspect the same is true of the second mechanism – that it depends on (or is greatly augmented by) the ability of a single mated wildtype female to give rise to brothers and sisters that can mate and establish a new population. This issue could be investigated by modifying our model to have local (as opposed to global) dispersal. In the continuous-space local-dispersal models of Champer *et al.* (2021), incorporating inbreeding depression appears to have the expected effect of increasing the likelihood of elimination.

The models presented here incorporate the stochastic effects that necessarily arise in dealing with discrete individuals, particularly at low densities, but nevertheless are explicitly solvable, requiring no stochastic simulations or generation of random numbers. They are also relatively simple, with only three parameters (*R*, *α*, *m* or *d*), and are not intended to give precise quantitative predictions about the consequences of a specific release in a specific species. Some of the previous simulation models that have shown population persistence have been substantially more complex, aiming to capture more faithfully the biology of one potential target species, *Anopheles gambiae*, the main vector of malaria in Africa (North *et al.*, 2013, 2019, 2020; Eckhoff *et al.*, 2017). Whether population persistence in these models is due solely to refugia and low density inbreeding, or whether some other features of the models (e.g., spatial and temporal heterogeneity, overlapping generations, etc) also promote population persistence remains to be determined. It would be interesting to include inbreeding depression or other strong Allee effects in these models to see how they affect the dynamics. There is good evidence of inbreeding depression in mosquitoes including *An. gambiae* (Armbruster *et al.*, 2000; Baeshen *et al.*, 2014; Turissini *et al.*, 2014; Ross *et al.*, 2019). We have also modelled inbreeding depression in a simple way, with only a single fixed fitness cost for females that have mated to a sibling, whereas it would be more realistic to have the costs increase with successive generations of inbreeding, or to explicitly model the deleterious recessive mutations that underlie inbreeding depression (Tanaka, 2000; Wittmann *et al.*, 2018). Over longer time periods these deleterious recessive mutations might get purged (Bundgaard *et al.*, 2021; Perez-Pereira *et al.*, 2021), though only if the population is not eliminated first.

## 6 Acknowledgements

This work benefited from useful discussions with Andrea Beaghton and Vasso Koufopanou, and was supported by the Bill & Melinda Gates Foundation (Grant number INV006610) and the Open Philanthropy Project (Grant number 2016-161185).

## 7 Supplementary Information

### 7.1 Scenario 6: Local reproduction followed by male dispersal and local mating

This scenario (and the one that follows) are hybrids of previous scenaria. Here, mated females settle randomly in patches, reproduce in a (locally) density dependent manner (as in Scenario 5) but instead of allowing local male and female progeny to mate, the male progeny from each patch migrate to the cloud and are then randomly re-assorted back to patches (whereas the female progeny remain in the patches they were born in). Mating then occurs locally (as in Scenario 5) and the resulting mated females rise back to the cloud to be re-assorted back to patches.

Similarly to previous scenaria, the cloud densities of W- and D-mated females in generation *t* are 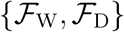, so any volume *V* in the cloud contains *on average* 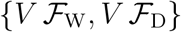 numbers of individuals, We again let the mated females in the cloud settle into patches with *average* numbers of individuals equal to 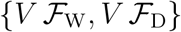. A random sample of {F_W_, F_D_} W- and D-mated females, different for each patch and Poisson-distributed with means 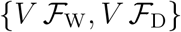, is drawn from the cloud and inserted in each patch.

As this scenario shares the same (local) density dependent reproduction mechanism with Scenaria 3 and 5, we can use (12) and (13) for the densities 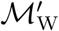 and 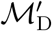 of the male offspring that migrate to the cloud. A random sample of {M_W_, M_D_} W- and D-males, different for each patch and Poisson-distributed with means 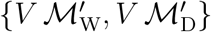, is then drawn from the cloud and inserted in each patch.

We use (9) for the probability ℙ (*i*_F_, *i*_W_, *j* – *i*_F_ – *i*_W_ | *j*, F_W_, F_D_) of having had {*i*_F_, *i*_W_, *j* – *i*_F_ – *i*_W_} female, W-male and D-male offspring in the patch (conditional onj total offspring from {F_W_, F_D_} mated females). However, the male offspring in the patch have now been replaced by {M_W_, M_D_} males that have settled in from the cloud.

The probability of *k* females, out of the *i*_F_ females in a patch, becoming W-mated females is 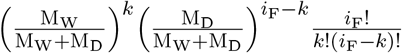 since every female undergoes a Bernoulli trial in picking a male out of the M_W_ W-males and M_D_ D-males in her patch. The expected number of W-mated females in the patch, conditional on {*i*_F_, M_W_, M_D_}, is thus

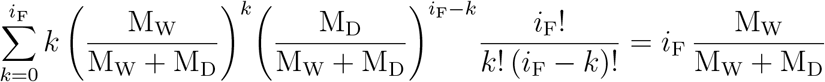

and, correspondingly, the expected number of D-mated females in the patch, conditional on {*i*_F_, M_W_, M_D_}, is 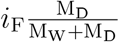.

We now derive the expected number of W- mated females in the patch, conditional on *i*_F_ local female offspring, by averaging over all possible Poisson-distributed M_W_ W- and M_D_ D-males in the patch (with means 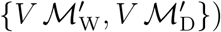) and using (12) and (13) for the densities 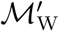 and 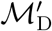:

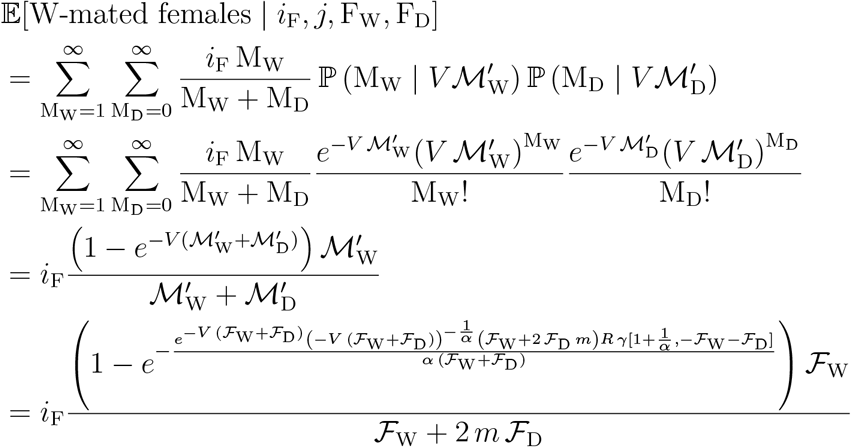

Similarly, the expected number of D-mated females in the patch, conditional on *i*_F_ local female offspring, is given by

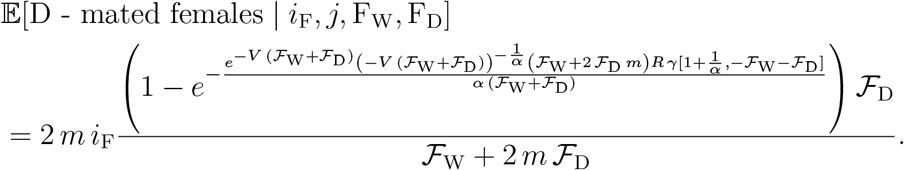

To obtain the expected number of W-mated females in the patch, conditional on {*j*, F_W_, F_D_}, we now sum over all possible *i*_F_ and *i*_W_:

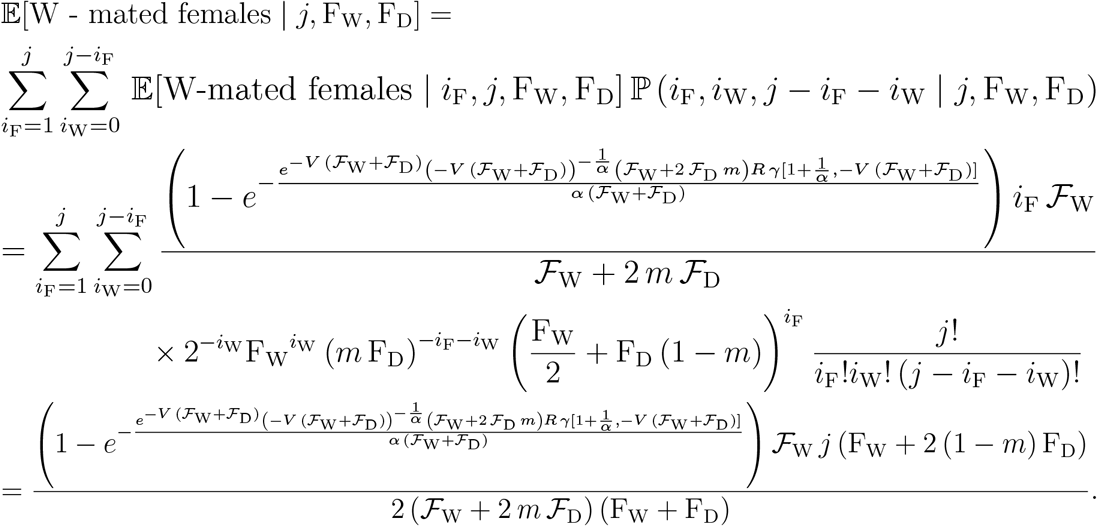

Similar analysis gives the expected number of D-mated females in each patch, conditional on {*j*, F_W_, F_D_}:

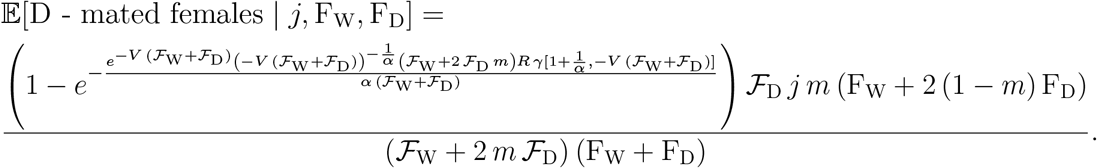

To calculate 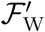, the density of W-mated females in the cloud in generation *t* + 1, we average the expected number of W-mated females in the patches conditional on {*j*, F_W_, F_D_} over all possible values of F_W_, F_D_ and *j* and divide by *V* to convert the mean number across all patches to a density in the cloud:

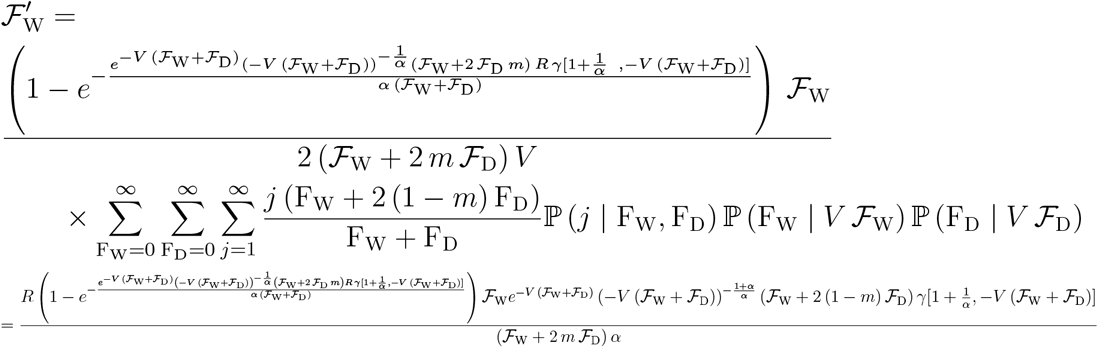

Similarly, 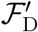, the density of D-mated females in the cloud in generation *t* + 1 is given by

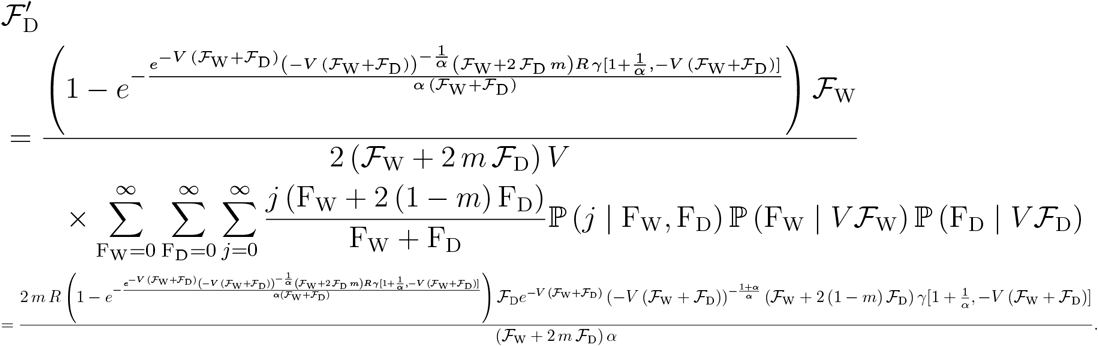

The change of variables 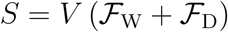 and 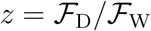 gives

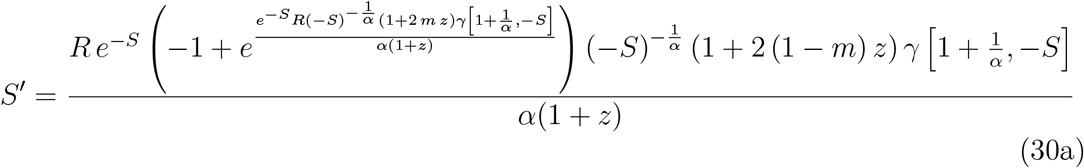

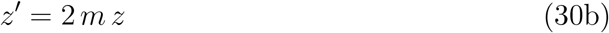

As in earlier scenaria, given that 2 *m* > 1, (30b) implies that *z* → ∞ as *t* → ∞ and the Driving Y asymptotically fixes in the population and, for sufficiently large *m*, the population will be eliminated.

### 7.2 Scenario 7: Local reproduction followed by female dispersal and local mating

In this scenario, mated females settle randomly in patches, reproduce in a (locally) density dependent manner (as in Scenario 5 and 6) but instead of allowing local male and female progeny to mate, the female progeny from each patch migrate to the cloud and are then randomly re-assorted back to patches (whereas the male progeny remain in the patches they were born in). Mating then occurs locally (as in Scenario 5 and 6) and the resulting mated females rise back to the cloud to be re-assorted back to patches.

Similarly to previous scenaria, the cloud densities of W- and D-mated females in generation *t* are 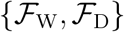, so any volume *V* in the cloud contains *on average* 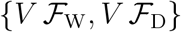 numbers of individuals, We again let the mated females in the cloud settle into patches with *average* numbers of individuals equal to 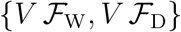. A random sample of {F_W_, F_D_} W- and D-mated females, different for each patch and Poisson-distributed with means 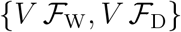, is drawn from the cloud and inserted in each patch.

As this scenario shares the same (local) density dependent reproduction mechanism with Scenaria 3, 5 and 6, we can use (11) for the densities 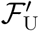 of the female offspring in the cloud. A random sample of F_U_ females, different for each patch and Poisson-distributed with mean *V* 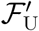, is drawn from the cloud and inserted in each patch.

We use (9) for the probability ℙ (*i*_F_, *i*_W_, *j* – *i*_F_ – *i*_W_ | *j*, F_W_, F_D_) of having had {*i*_F_, *i*_W_, *j* – *i*_F_ – *i*_W_} female, W-male and D-male offspring in the patch (conditional on *j* total offspring from {F_W_, F_D_} mated females). However, the *i*_F_ female offspring in the patch have now been replaced by F_U_ females that have settled in from the cloud.

The probability of *k* females, out of the F_U_ females that settle in a patch, becoming W-mated females is 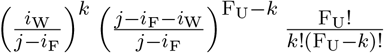 since every female undergoes a Bernoulli trial in picking a male out of the *i*_W_ W-males and *i*_D_ D-males in her patch. The expected number of W-mated females in the patch, conditional on {F_U_, *i*_W_, *j* – *i*_F_ – *i*_W_}, is thus

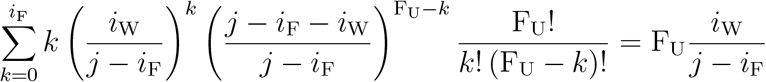

and, correspondingly, the expected number of D-mated females in the patch, conditional on {F_U_, *i*_W_, *j* – *i*_F_ – *i*_W_}, is 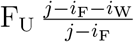

To obtain the expected number of W-mated females in the patch, conditional on {*j*, F_W_, F_D_}, we now sum over all possible *i*_F_ and *i*_W_ (note that *i*_F_ = *j* is excluded from the *i*_F_-summation as it would imply no male offspring for the females to mate with), noting that the W-male fraction is 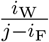:

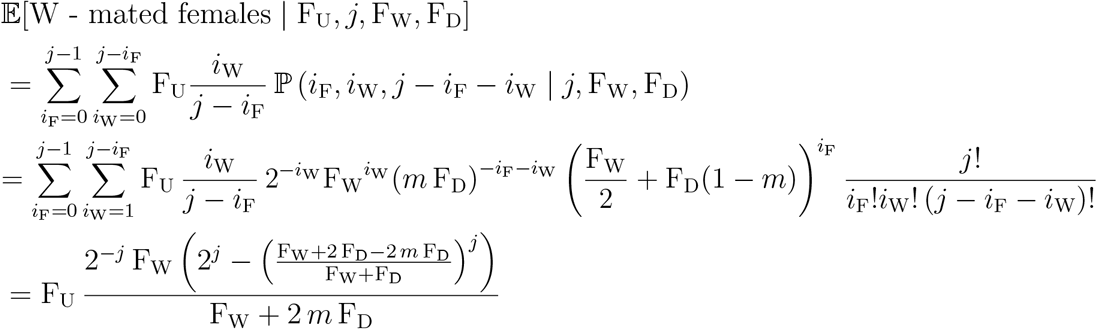

We now sum over all possible values of F_U_ females:

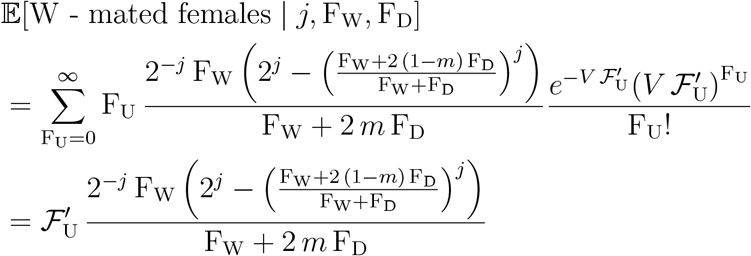

To calculate 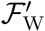, the density of W-mated females in the cloud in generation *t* + 1, we sum over all possible values of F_W_, F_D_ and *j* and divide by *V*; we use (11) for the cloud density 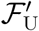 of unmated females that settle in the patches before mating:

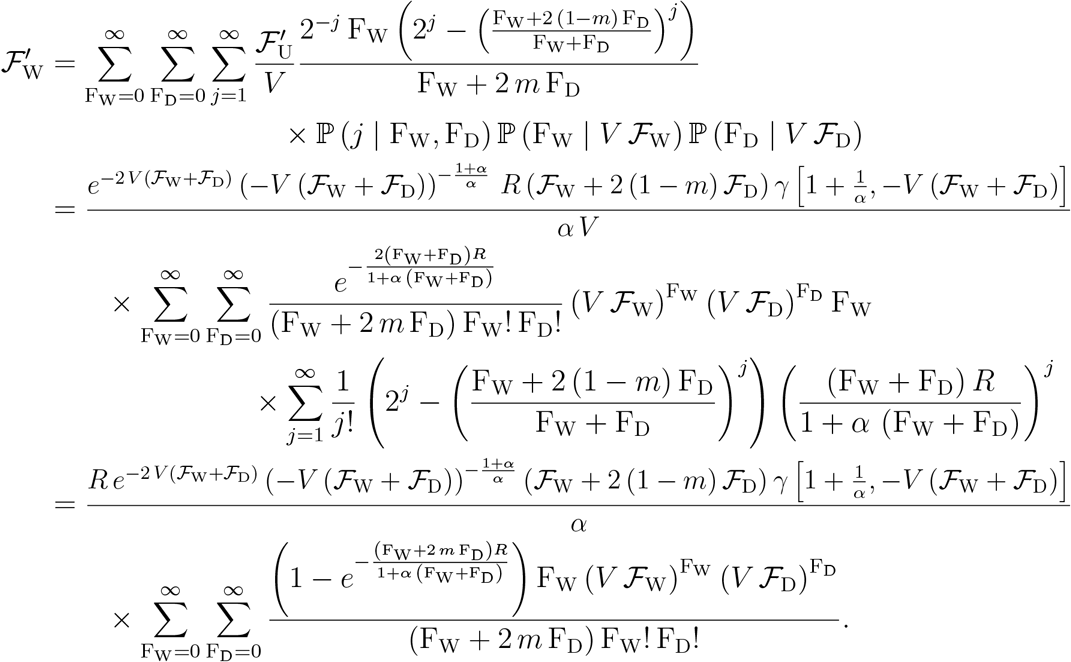

Similarly, we obtain 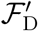, the density of D-mated females in the cloud in generation *t* + 1:

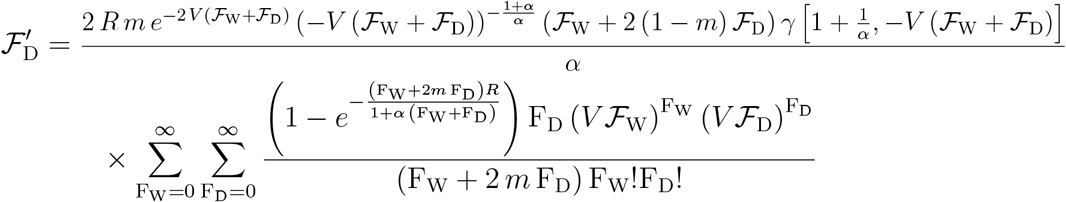

The change of variables 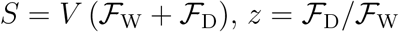, and F = F_W_ + F_D_ gives

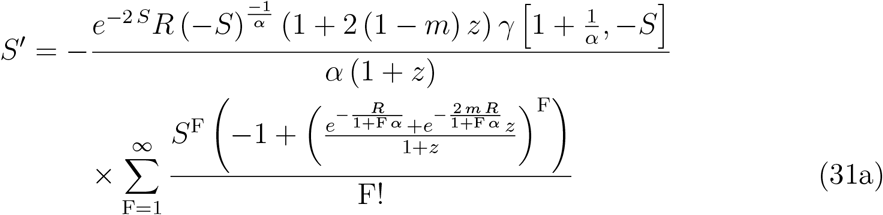

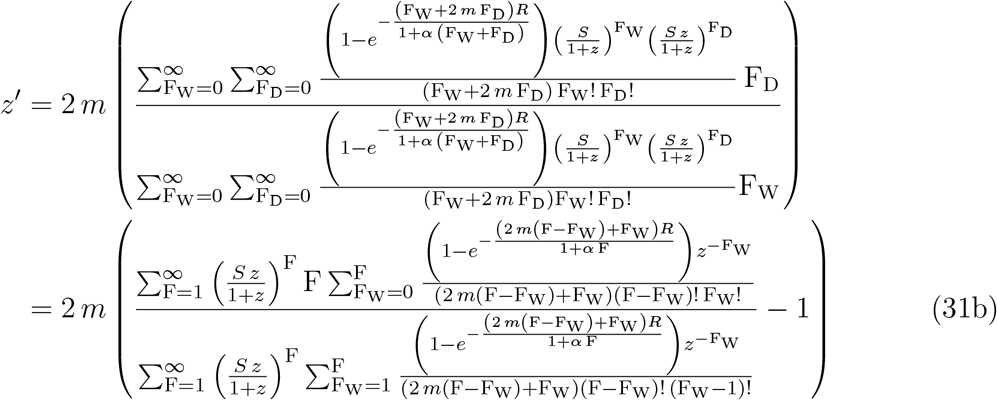

Numerical solution of (31a)-(31b) for a wide range of parameters and initial conditions indicates that *z* → ∞ as *t* → ∞ and the Driving Y asymptotically fixes in the population and, for sufficiently large *m*, the population will be eliminated.

### 7.3 Proof used in Section 4.1

In order to remove, on average, a fraction 0 ≤ *δ* ≤ 1 of sibling-mated females from the total number of new mated females in the patch, we first prove that the expected number of new sibling-mated W-mated females in the patch, conditional on 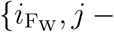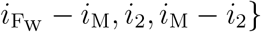 offspring from {F_W_, F_D_} mated females, is 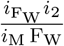. A similar proof, not shown here, gives the expected number of new sibling-mated D-mated females in a patch, conditional on 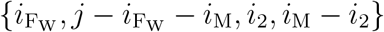 offspring from {F_W_, F_D_} mated females, as 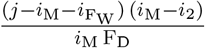.

*Proof.* We assume that there are F_W_ and F_D_ W- and D-mated females, respectively, in a patch. The W-mated females produce 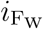 female and *i*_2_ W-male offspring. We will prove by mathematical induction that the expected number of female offspring that will mate with sibling W-males in the patch is 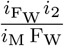, where *i*_M_ is the total number of male offspring in the patch (i.e. the sum of W- and D-males):

Step 1 – prove it is valid for F_W_ = 2:

We set F_W_ = 2 and F_D_ arbitrary. Let W-mated female #1 produce *x*_1_ females and *y*_1_ W-males and W-mated female #2 produce 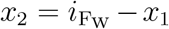 females and *y*_2_ = *i*_2_ – *y*_1_ W-males. The expected number of new W-mated females that have mated with siblings (conditional on {*x*_1_, *y*_1_}) is 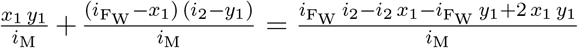. The probability of female #1 having *x*_1_ female and *y*_1_ W-male offspring is 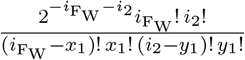, so the expected number of W-mated females from sibling pairings is:

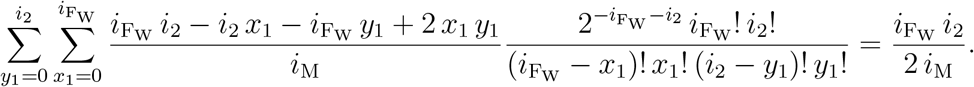

Step 2 – assume it is valid for arbitrary F_W_ > 2:

It is assumed that the expected number of W-mated females from sibling pairings between offspring of the F_W_ W-mated mothers in the patch is 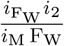.

Step 3 – prove it is valid for F_W_ + 1:

We divide the offspring of F_W_ + 1 W-mated females in one cohort from F_W_ mothers and another one from the (F_W_ + 1)-th mother. The first cohort contains 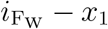 and *i*_2_ – *y*_1_ offspring and the 2nd cohort contains *x*_1_ and *y*_1_ offspring. The probability of the 2^nd^ cohort containing *x*_1_ and *y*_1_ offspring is 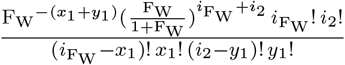. The expected number of new W-mated females that have mated with siblings (conditional on {*x*_1_, *y*_1_}) is 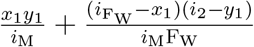 (the division by F_W_ in the 2^nd^ fraction is a direct consequence of our assumption in Step 2, i.e. that, on average, a fraction 1/F_W_ of the total new W-mated females in the patch are sibling-mated). The expected number of W-mated females from sibling pairings is thus:

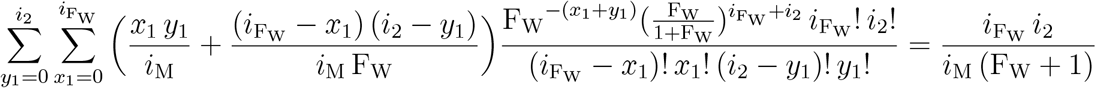

This completes the mathematical induction proof that the expected number of new W-mated females from sibling pairings, conditional on 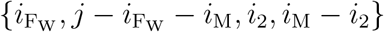 offspring from {F_W_, F_D_} mated females in a patch, is 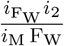.

